# Single cell eQTL analysis identifies cell type-specific genetic control of gene expression in fibroblasts and reprogrammed induced pluripotent stem cells

**DOI:** 10.1101/2020.06.21.163766

**Authors:** Drew Neavin, Quan Nguyen, Maciej S. Daniszewski, Helena H. Liang, Han Sheng Chiu, Anne Senabouth, Samuel W Lukowski, Duncan E. Crombie, Grace E. Lidgerwood, Damián Hernández, James C. Vickers, Anthony L. Cook, Nathan J. Palpant, Alice Pébay, Alex W. Hewitt, Joseph E. Powell

## Abstract

The discovery that somatic cells can be reprogrammed to induced pluripotent stem cells (iPSCs) - cells that can be differentiated into any cell type of the three germ layers - has provided a foundation for *in vitro* human disease modelling^1,2^, drug development^1,2^, and population genetics studies^3,4^. In the majority of instances, the expression levels of genes, plays a critical role in contributing to disease risk, or the ability to identify therapeutic targets. However, while the effect of the genetic background of cell lines has been shown to strongly influence gene expression, the effect has not been evaluated at the level of individual cells. Differences in the effect of genetic variation on the gene expression of different cell-types, would provide significant resolution for in vitro research using preprogramed cells. By bringing together single cell RNA sequencing^15–21^ and population genetics, we now have a framework in which to evaluate the cell-types specific effects of genetic variation on gene expression. Here, we performed single cell RNA-sequencing on 64,018 fibroblasts from 79 donors and we mapped expression quantitative trait loci (eQTL) at the level of individual cell types. We demonstrate that the large majority of eQTL detected in fibroblasts are specific to an individual sub-type of cells. To address if the allelic effects on gene expression are dynamic across cell reprogramming, we generated scRNA-seq data in 19,967 iPSCs from 31 reprogramed donor lines. We again identify highly cell type specific eQTL in iPSCs, and show that that the eQTL in fibroblasts are almost entirely disappear during reprogramming. This work provides an atlas of how genetic variation influences gene expression across cell subtypes, and provided evidence for patterns of genetic architecture that lead to cell-types specific eQTL effects.

## Introduction

Mapping expression quantitative trait loci (eQTL) is a powerful method to study how common genetic variation between individuals influences gene expression^5,6^. To date, nearly all eQTL studies have been conducted while interrogating ‘bulk’ samples, where the RNA is collected from millions of lysed cells, and therefore gene expression represents an average across all cells in a sample. However, even with ‘bulk’ RNA analyses, comparisons of eQTL identified from different tissues^7,8^, and cultured cell lines^9,10^ has revealed differences in both the presence, and the directions of allelic effects of eQTL^13,14^. In stem cell systems, single cell approaches have already revealed that cell cultures do not contain homogeneous cell types^60^, instead consisting of multiple cell types that have different transcriptional profiles. By harnessing technology and recent methods that enable high-throughput generation of single cell data using cell multiplexing across donors^22–24^, provides an experimental framework in which cell-type specific genetic effects on gene expression can be tested - permitting the identification of eQTL that are truly cell type specific, and that would otherwise be undetected by ‘bulk’ approaches.

Previous studies have identified cell type specific eQTL using scRNA-seq which were unobservable in bulk RNA-sequence studies^25–29^. The first study to report this enhanced cell type specific eQTL detection from scRNA-seq investigated 92 genes measured in 1,440 single cells from lymphoblastoid cell lines in 15 individuals^27^. In the current study, we set out to understand the impact of single nucleotide polymorphisms (SNPs) - common genetic variants - on gene expression in fibroblast and reprogrammed iPSC cell types through eQTL mapping at the level of cell subpopulations.

## Results

To identify cell-type specific eQTL in an unbiased manner, we generated scRNA-seq expression profiles of 83,985 cells - 64,018 cultured dermal fibroblasts, generated from skin biopsies from 79 unrelated individuals, and 19,967 iPSCs reprogrammed from 31 of the dermal fibroblast lines (**Figure 1A**). After quality control, we used an unbiased approach to map cells to reference transcriptomes from the human primary cell atlas^30,31^, demonstrating that the majority of fibroblasts mapped to the fibroblast reference, while the majority of iPSCs mapped to the iPSC or embryonic stem cell references. We used an unsupervised clustering approach^32^ to identify six types of fibroblasts and four types of iPSCs (**Figures 1B-C** and **S1**). Fibroblast and iPSC types contained equal distributions of individual donors, pool batches and cell cycle states (**Figures S2 and S3**). Cell types of fibroblasts and iPSCs were classified based on the expression of key marker genes. For fibroblasts: TUBA1B^hi^/DCN^lo^; CD9^hi^/FTL^lo^; CD9^hi^/C1R^hi^; DCN^hi^/C1R^hi^; WISP2^hi^/THBS1^hi^; and TUBA1B^lo^/CD9^lo^. And the iPSCs: FTL^hi^/ENO1^hi^; FTL^lo^/BST2^hi^; FTL^lo^/SNHG8^hi^; and FTL^hi^/ENO1^lo^. (**Figures 1D-E, S4** and **S5 and Tables S1 and S2**). Further, pseudo-trajectory analysis demonstrated that the identified cell types were positions along a clear lineage trajectory for both fibroblast and iPSC types (**Figure S6**).

**Figure 1:**
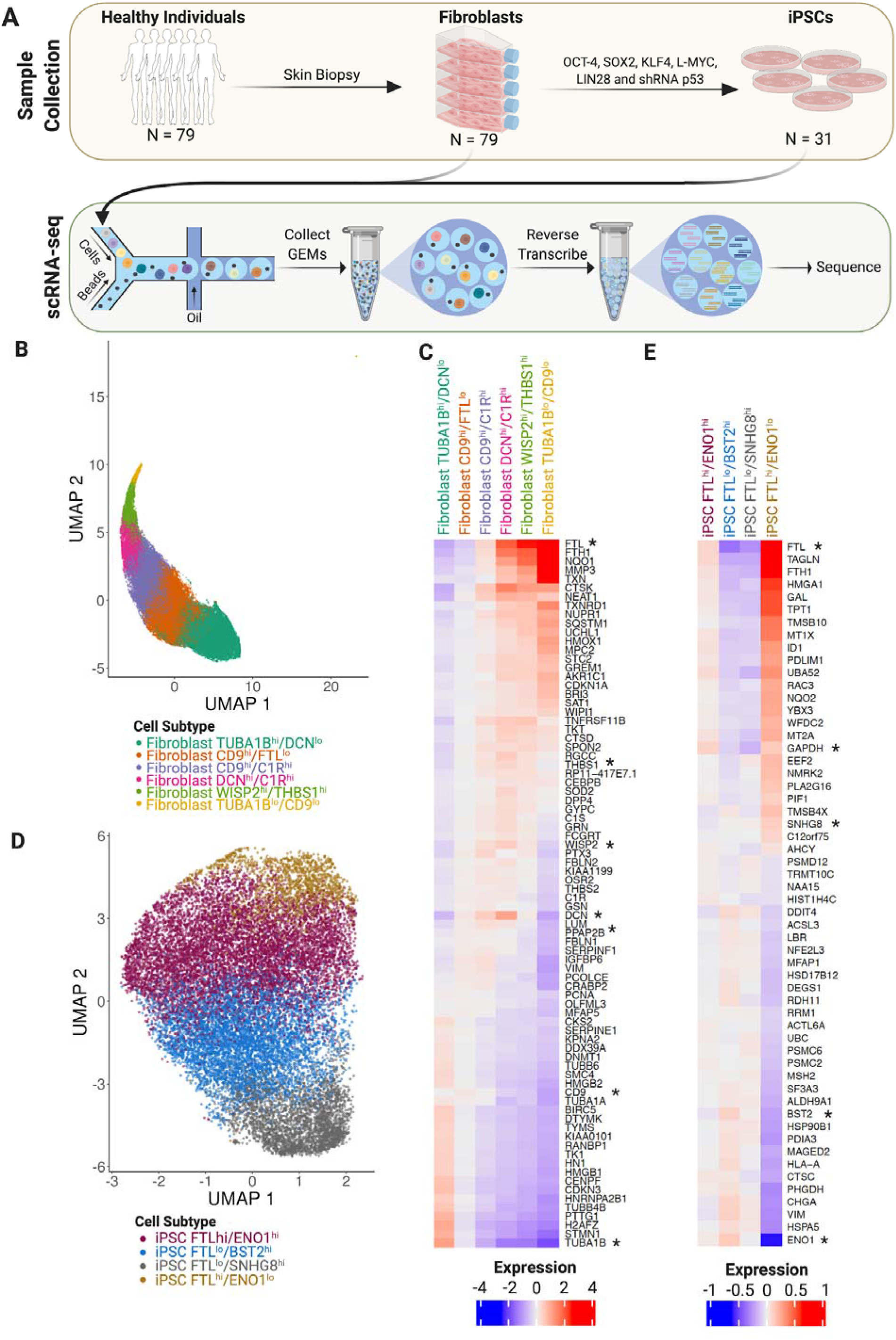
Fibroblast and iPSC Cluster Characterization. **A)** This study used skin biopsies to generate fibroblasts from 79 healthy volunteers and reprogrammed them into induced pluripotent stem cell (iPSC) lines for 31 of the original 79 individuals. **B)** Six fibroblast subtypes were identified from the transcriptional profiles of 64,018 single fibroblast cells. **C)** The top 20 differentially expressed genes from each fibroblast subtype demonstrate a continuum of expression across the six fibroblast subtypes. **D)** Four iPSC subtypes were identified from 19,967 single iPSCs. **E)** The top 20 differentially expressed genes from each iPSC subtype. *Indicates the genes used to name each subtype.

We subsequently tested for cis-eQTL independently in each of the 10 cell types. We identified a total of 30,574 eQTL for 1,951 genes (FDR < 0.05) across all cell types - 29,800 eQTL for 1,877 genes in fibroblast types and 774 cis-eQTL for 85 genes in iPSC types (**Table S3 and S4**). Assessing the overlap of eQTL and eGenes, revealed that the majority of cis-eQTL are predominantly cell type specific, with 82.4% of the eGenes (65.4% of the cis-eQTL) identified in only one fibroblast type (**Figures 2A-B** and **S7A**) and 97.6% of the eGenes (99.6% of the cis-eQTL) identified in only one iPSC type (**Figure S7B-C**). Cell-type ubiquitous (shared across cell sub-types) were rare, with eight eGenes with eQTL in all fibroblast types (**Figures 2A, S7A, S8**), and none across all iPSC types (**Figure S7B**). Looking across the cell reprogramming event, we observed a complete lack of shared eQTL between fibroblast and iPSCs. Only 11 genes had eQTL in both fibroblasts and iPSCs (Figure S7E), but none of those shared a common eSNP, or SNPs in linkage disequilibrium with one another (r >0.2), indicating that their expression was likely associated with independent loci (**Figures 2C, S9**).

**Figure 2:**
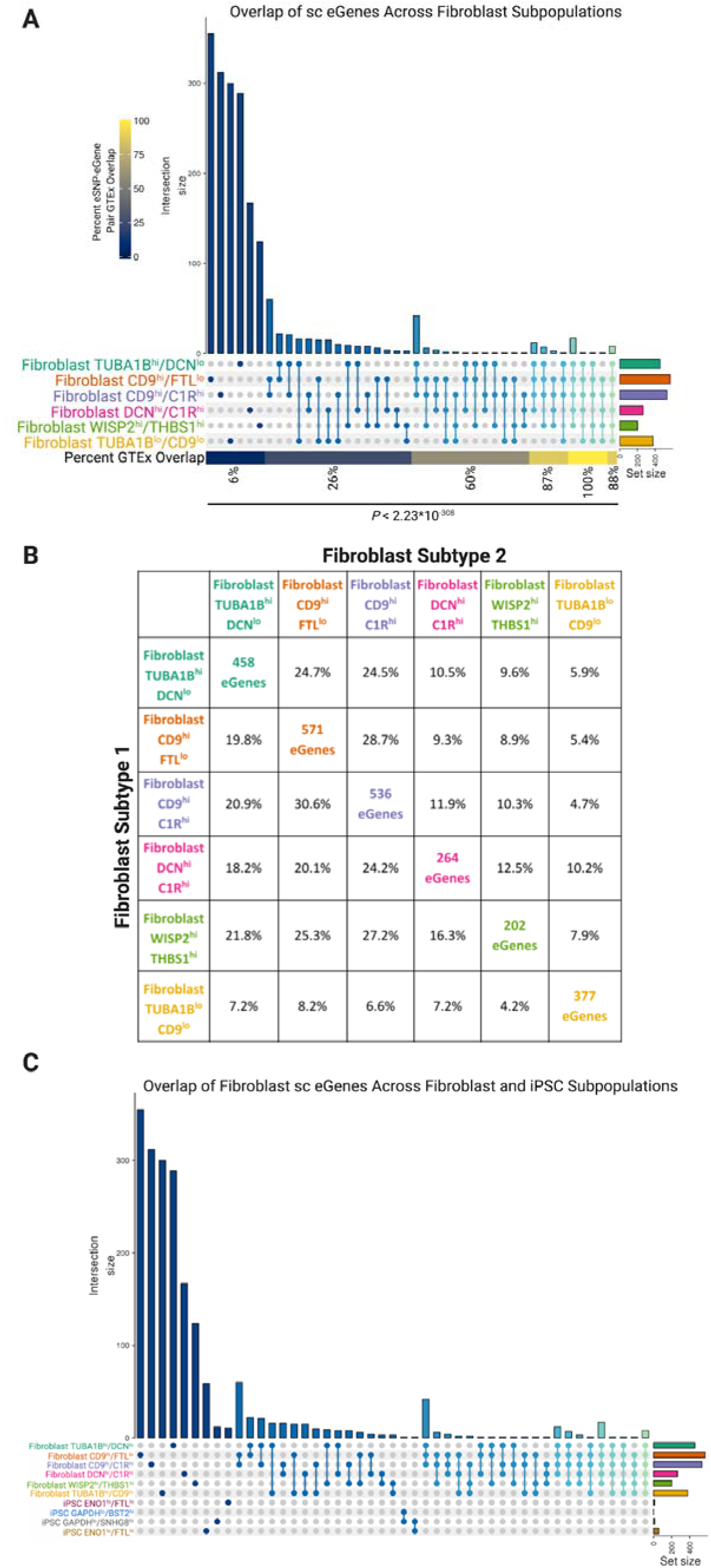
Identification of single cell eQTL in fibroblast and iPSC subtypes. **A)** The majority of single cell (sc) eGenes in fibroblasts are subtype-specific. Further, the single cell eGenes that were detected in two or more fibroblast subtypes were significantly more likely to be detected as eQTL in bulk fibroblast RNA-sequence data from the gene tissue expression (GTEx) database (*P*=0, Cochran-Armitage Test). **B)** The total number of eGenes and percent that are also observed in other fibroblast subtypes further shows that most eGenes are unique to a given subtype. **C)** None of the single cell eGenes-eSNP pairs that were observed in fibroblasts were observed in the iPSC subtypes that were generated from the same individuals.

We then investigated whether the eQTL identified in fibroblasts replicated in bulk RNA-sequence data from the Genotype-Tissue Expression (GTEx, culture fibroblasts n=483)^33^. Only 12% of the 29,800 eQTL identified in the six fibroblast types replicated (p<0.05/29,800) in GTEx, although they had a consistent shared direction of allelic effects. Given the high percentage of cell-type specific eQTL, one explanation for this observation is that bulk RNA approaches mask cell-type specific effects through averaged gene expression across cells. From this, we hypothesised that cell-type ubiquitous eQTL (from the single cell analysis) would have higher replication rates compared to cell-type specific eQTL. Testing for replication for eQTL that were shared across multiple fibroblast cell types in the scRNA-seq, showed a highly significant difference compared with eQTLs that were significant in just one fibroblast type (p<2^−308^ for eGenes and p=1^−108^ for eSNPs; **Figures 2A and S7A**). Further, we identified that the allelic effect size of the eGenes and eQTLs in GTEx cultured fibroblasts was positively correlated with the number of fibroblast types where those eGenes and eQTL were significant (**Figure S10**). These results indicate that eQTL mapping using bulk RNA-sequence data is likely not sensitive enough to identify fibroblast type-specific eQTL.

Based on our initial observation of the specificity of cell type eQTL effects, we next sought to identify how different types of genetic architecture and gene expression patterns contributed to the cell-type specific effects in fibroblasts and iPSCs.

One potential explanation for the cell type-specific eQTL detection, is that the gene is only expressed in one cell type, and therefore, we would not expect to observe an eQTL in the other cell types that where the gene isn’t expressed. To evaluate this, we correlated the expression of each gene that had a significant cell-type specific eQTL effect, with its expression levels in each of the other cell types. (**Figure S11, S12**). These results indicate that cell type-specific eQTL are not a function of cell type-specific gene expression, showing high levels of correlation in almost all instances. Another possible explanation for the cell type-specific eQTL is low statistical power to detect eQTL in multiple cell types. To assess this hypothesis, we implemented an empirical framework to test the rank of the test statistics for eGene SNP effects across the non-significant cell types for each cell-type specific eQTL. In almost all instances we observed none, or very limited enrichment of the test statistic across cell types (**Figure S12**). In the instance where we identified significant enrichment, it existed between the CD9^hi^/FTL^lo^ and CD9^hi^/CLR^hi^ fibroblast cell-types that are similar to one another (**Figure 1**). Therefore, we conclude that the majority of cell type-specific eQTL that we have identified were not a result of differences in gene expression or due to lack of statistical power. We next set out to interrogate eGenes that were in common between multiple cell types.

We identified 283 eGenes that were significant in multiple cell types, but which had different top eSNPs - 255 eGenes in at least two fibroblast types, no eGenes with different top eSNPs in two iPSC types and 11 eGenes with different top eSNPs in a fibroblast type and an iPSC type. In these instances, we considered two alternative hypotheses: 1) that there was one eQTL shared between cell types but that it was tagged by a different top eSNP in each cell type, or 2) that there were two independent cell type-specific eQTLs for the same gene. To address these hypotheses, we tested whether the top eSNP in a given cell type was still significantly associated with gene expression after correcting for the top eSNP in the other cell type. A significant association of the SNP with the eGene expression after correction for the other eSNP would indicate that the two eSNPs were not tagging the same eQTL and were, therefore, independent loci. The analysis identified that between 44.4% and 73.8% of these loci for a given fibroblast type were independent (**Figure 3 and Table S5**), and 100% of the eGenes shared between the fibroblast and iPSC types were also independent loci (**Table S6**). These results denote that many of the eGenes that were shared between multiple cell types, are in fact regulated by different loci, providing further support to our previous finding that the majority of eQTL are cell type-specific.

**Figure 3:**
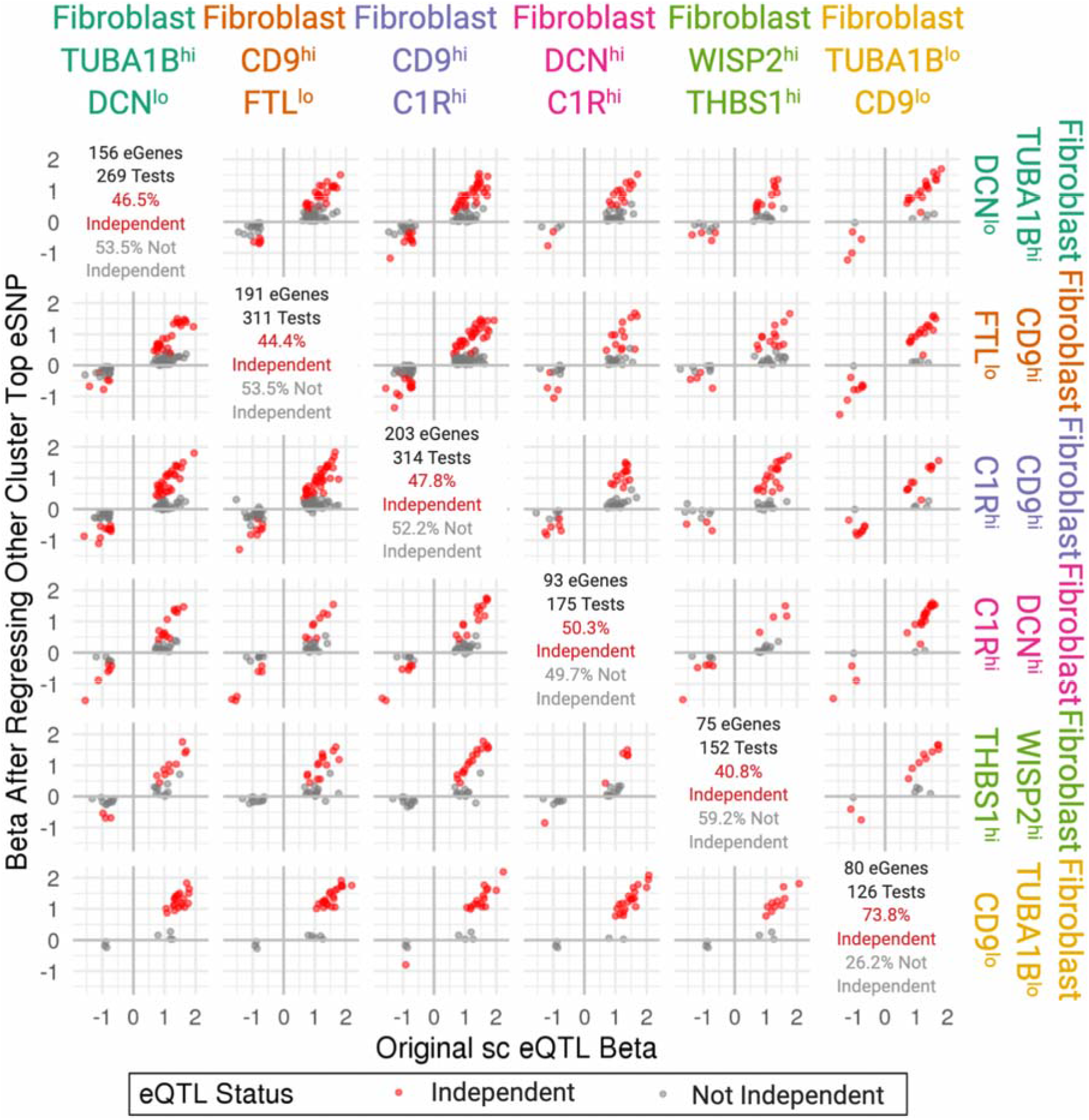
eGene Comparison across Fibroblast subtypes. **A)** The correlation of fibroblast eGenes from subtype 1 (x-axis) with fibroblast eGenes from subtype 2 (y-axis) demonstrates that eGenes are similarly expressed across all fibroblast subtypes. **B)** eGenes that were shared between at least two fibroblast subtypes were tested for independence. The top eSNP for eGenes that were shared between two fibroblast subtypes was regressed from the other subtype in order to test if those were independent eSNP loci. Many (40-73%) of the fibroblast top eSNPs remained significant after regression of the top eSNP from another fibroblast subtype.

Next, we investigated the 153 eGenes that shared at least one significant eSNP-eGene pair (i.e. same top eSNP) across multiple fibroblast types. We evaluated the potential interactions between cell type and eSNP, leading to difference magnitude of the allelic effect in different cell types by testing for a SNP-fibroblast cell type interaction for each of the 153 eSNP-eGene pairs. In cases where multiple eSNP-eGene pairs were significant for the same eGene across multiple cell types, we tested the eSNP-eGene pair with the largest beta difference between two fibroblast types. This analysis identified 64 (41.8%) significant eSNP-fibroblast type interactions (**Table S7**). This analysis identifies instances where there are eQTL that are ubiquitous, but whose alleleic effect significantly varies across cell types.

After identifying that the majority of eQTL are cell type-specific, and that the cell type can interact with the SNP locus to alter the allelic effect, we interrogated our results for loci that were statistically significant across multiple analyses. We first asked whether any eGene was significant in all six fibroblast types and was also a significant interaction between cell type and eQTL, and identified the guanylate binding protein 3 (GBP3) locus that was significant in both tests (**Figure 4A-B**). The rs541032500-GBP3 locus has a significant eQTL in five of the six fibroblast cell types, and demonstrated a significant interaction with the cell type (p=1.5^−02^). Interestingly, GBP3 has been shown to be induced in fibroblasts by interferon treatment^34,35^. Our results indicate that the effect of interferon induction of GBP3 is likely to be mediated by the genotypes carries at the rs541032500 loci, and it’s magnitude vary based on the cell type context.

**Figure 4:**
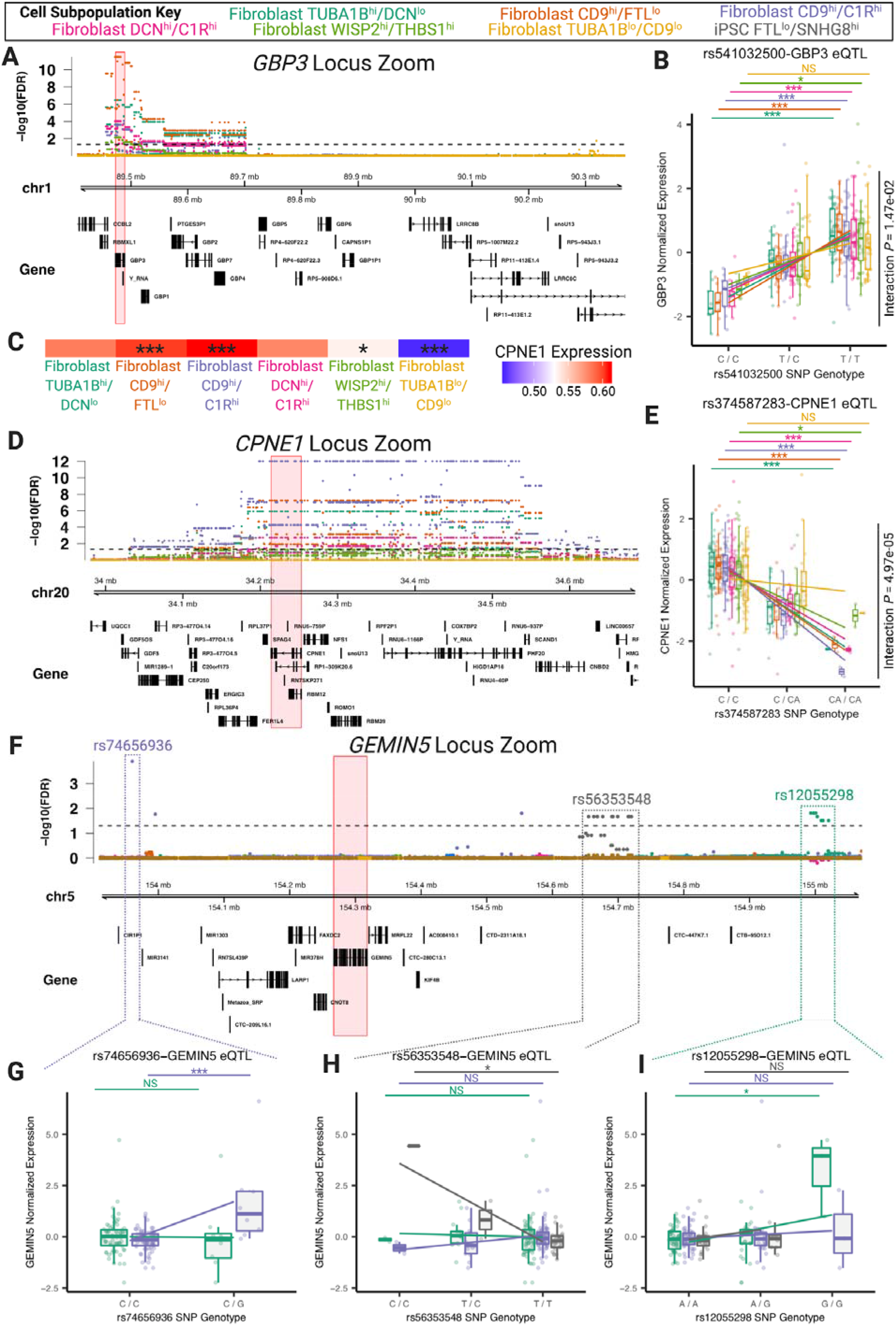
Examples of eQTL identified in fibroblast and iPSC subtypes. **A)** The *GBP3* gene was the only eGene that was significant in all fibroblast subtypes and also **B)** demonstrated an interaction between the fibroblast subtypes and the eQTL. **C)** CPNE1 was differentially expressed across the fibroblast subtypes and **D)** was a significant eGene in five of the six fibroblast subtypes. **E)** Further, the rs3474587283-*CPNE1* eQTL demonstrated striking subtype by SNP interaction. **E)** The *GEMIN5* gene was an eGene in two fibroblast subtypes and one iPSC subtype the top eSNPs were different in each subtype. The CD9^hi^/C1R^hi^ fibroblast subtype eQTL was 3’ of the GEMIN5 gene, the GAPDH^hi^/SNHG8^hi^ iPSC subtype eQTL was 5’ of the *GEMIN5* gene and the TUBA1B^hi^/DCN^lo^ fibroblast subtype was further 5’ of the *GEMIN5* gene. All three loci were independent of one another (*P* < 0.05). **F)** The top eSNP (rs74656936) for the *GEMIN5* gene in the CD9^hi^/C1R^h^ fibroblast subtype was not significant in the TUBA1B^hi^/DCN^lo^ fibroblast subtype and the rs74656936 was not frequent enough in the iPSC lines to be analyzed. **G)** The top eSNP (rs56353548) for the *GEMIN5* gene in the GAPDH^hi^/SNHG8^hi^ iPSC subtype was not significant in either the CD9^hi^/C1R^hi^ fibroblast subtype or the TUBA1B^hi^/DCN^lo^ fibroblast subtype. **H)** The top eSNP (rs12055298) for the *GEMIN5* gene in the TUBA1B^hi^/DCN^lo^ fibroblast subtype was not significant in the CD9^hi^/C1R^h^ fibroblast subtype or the GAPDH^hi^/SNHG8^hi^ iPSC subtype. \**P* < 0.05; \*\**P* < 0.01; \*\*\**P* < 0.001; NS=non-significant.

Following this, we evaluated whether any of the differentially expressed genes (**Figures 1C, S4 and Table S1**) also had a significant interaction with eQTL. From this we identified the copine 1 (CPNE1) locus, which was significantly increased in the CD9^hi^/FTL^lo^ and the CD9^lo^/C1R^hi^ fibroblast types, and significantly decreased in the WISP2^lo^/THBS1^hi^ and TUBA1B^lo^/CD9^lo^ fibroblast types (**Figure 4C**). In addition, the rs374587283-CPNE1 cis-eQTL was significant in five of the six fibroblast types (**Figure 4D-E**) and demonstrated a significant SNP-cell type interaction (p=4.9^−05^; **Figure 4E**).

Finally, we investigated whether any of the eGenes had evidence for association with multiple independent loci using conditional association analysis (Methods). One example of this genetic architecture is the gem nuclear organelle associated protein 5 (GEMIN5) locus, which has three independent loci in three cell types - two fibroblast types and one iPSC type (**Figure 4F-I**). GEMIN5 was a significant eGene for the TUBA1B^hi^/DCN^lo^ fibroblast type, the CD9^hi^/C1R^hi^ fibroblast type and the FTL^lo^/SNHG8^hi^ iPSC type. However, the top eSNP from each cell type for GEMIN5 are independent of one another. For example, the rs74656936-GEMIN5 eQTL was significant in the CD9^hi^/C1R^hi^ but not the TUBA1B^hi^/DCN^lo^ fibroblast cell types (**Figure 4G**). Meanwhile, the rs5635348-GEMIN5 eQTL was significant in the FTL^lo^/SNHG8^hi^ iPSC type but was not significant in either of the fibroblast types (**Figure 4H**). Finally, the rs12055298-GEMIN5 eQTL was significant in the TUBA1B^hi^/DCN^lo^ fibroblast type but not in the CD9^hi^/C1R^hi^ fibroblast type or the FTL^lo^/SNHG8^hi^ iPSC type (**Figure 4I**).

We set out to identify and define the dynamics of eQTL in fibroblasts and fibroblast-derived iPSC cell types. Collectively, our results provide evidence that there is a high degree of cell type-specific gene regulation that is not captured with bulk RNA-seq. Further, our results indicate that even when the same eGene is observed in different cell types, the allelic effect may be altered in different cell types, or may be regulated by different loci entirely. Our findings support previous reports that many cell type-specific eQTL are not detected using bulk RNA-sequencing and that scRNA-seq can be utilised to enhance eQTL detection^36^.

scRNA-seq provides a number of advantages over bulk RNA-sequencing for eQTL mapping. Specifically, scRNA-seq enables cell types to be identified in an unbiased manner before eQTL detection. Therefore, even cell types that have previously not been described or well characterised can be identified and separated for eQTL mapping, thereby decreasing the measurement noise that is introduced due to heterogeneity of cells in bulk RNA-sequence profiling. Furthermore, scRNA-seq enables the cells from multiple individuals to be pooled in a single experiment, thereby decreasing technical batch effects that can confound biological variation between individuals. Finally, this study has provided a map of eQTL in fibroblast and fibroblast-derived iPSC types that will be an important reference for future studies in iPSC-derived cell types.

## Methods

### Participant Recruitment and Ethics Approval

Experimental work was approved by the Human Research Ethics committees of the Royal Victorian Eye and Ear Hospital (11/1031), University of Melbourne (1545394), University of Tasmania (H0014124) in accordance with the requirements of the National Health & Medical Research Council of Australia (NHMRC) and conformed with the Declaration of Helsinki^37^.

### Fibroblast culture

Human skin punch biopsies were obtained from subjects over the age of 18 years. Fibroblasts were cultured in DMEM high glucose supplemented with 10% foetal bovine serum (FBS), L-glutamine, penicillin (100 U/mL), streptomycin 100 (µg/mL) (all from Thermo Fisher Scientific, USA). All cell lines were mycoplasma-free (MycoAlert mycoplasma detection kit, Lonza, Switzerland).

### Generation and maintenance of iPSCs

Human iPSCs were reprogrammed from fibroblast cultures by nucleofection (Amaxa™ Nucleofector™) of episomal vectors expressing OCT-4, SOX2, KLF4, L-MYC, LIN28 and shRNA against p53^38^, in feeder- and serum-free conditions using TeSR™-E7™ medium (STEMCELL Technologies, Canada) and selected by sorting with anti-human TRA-1-60 Microbeads using a MultiMACS (Miltenyi, Germany) as described by Crombie et al^39^ and Daniszewski et al^40^. Cells were maintained on vitronectin XF™ (STEMCELL Technologies™) -coated plates using TeSR™-E8™ (Stem Cell Technologies). At passage eight, cells were assessed for quality control as described previously^40^.

### iPSC quality control

Pluripotency was assessed by immunochemistry for expression of OCT3/4 (sc-5279, Santa Cruz Biotechnology, USA) and TRA-1-60 (MA1-023-PE, Thermo Fisher Scientific). Copy number variation (CNV) analysis of original fibroblasts and iPSCs was performed using Illumina HumanCore Beadchip arrays with PennCNV^41,42^ and QuantiSNP^42^ with default parameter settings. Chromosomal aberrations were deemed to involve ≥ 20 contiguous SNPs or a genomic region spanning ≥ 1MB ^41,42^. The B allele frequency (BAF) and the log R ratio (LRR) were extracted from GenomeStudio (Illumina, USA) for representation.

### Generating the single cell RNA-sequence data

Viable cells were sorted on a BD Influx cell sorter (Becton-Dickinson) using Propidium Iodide into Dulbecco's phosphate buffered saline (PBS) + 0.1% bovine serum albumin and retained on ice. Sorted cells were counted and assessed for viability with Trypan Blue using a Countess automated counter (Invitrogen), and then resuspended at a concentration of 800-1000 cells/µL (8 × 10^5^ to 1 × 10^6^ cells/mL). Final cell viability estimates ranged between 92-96%.

Single cell suspensions were loaded onto 10X Genomics Single Cell 3' Chips along with the reverse transcription (RT) mastermix as per the manufacturer's protocol for the Chromium Single Cell 3' Library (10X Genomics; PN-120233), to generate single cell gel beads in emulsion (GEMs). Reverse transcription was performed using a C1000 Touch Thermal Cycler with a Deep Well Reaction Module (Bio-Rad) as follows: 55°C for 2h; 85°C for 5min; hold 4°C. cDNA was recovered and purified with DynaBeads MyOne Silane Beads (Thermo Fisher Scientific; Cat# 37002D) and SPRIselect beads (Beckman Coulter; Cat# B23318). Purified cDNA was amplified as follows: 98°C for 3min; 12x (98°C for 15s, 67°C for 20s, 72°C for 60s); 72°C for 60s; hold 4°C. Amplified cDNA was purified using SPRIselect beads and sheared to approximately 200bp with a Covaris S2 instrument (Covaris) using the manufacturer’s recommended parameters. Sequencing libraries were generated with unique sample indices (SI) for each chromium reaction. Libraries were multiplexed, and sequenced on an Illumina NextSeq 500 (NextSeq control software v2.0.2/Real Time Analysis v2.4.11) using a 150-cycle NextSeq 500/550 High Output Reagent Kit v2 (Illumina, FC-404-2002) in standalone mode as follows: 98 bp (Read 1), 14 bp (I7 Index), 8 bp (I5 Index), and 10 bp (Read 2).

### scRNA-seq Cellranger Processing

Processing of the sequencing data into transcript count tables was performed using the Cell Ranger Single Cell Software Suite by 10X Genomics (http://10xgenomics.com/). Raw base call files from the NextSeq 500 sequencer were demultiplexed, using the cellranger mkfastq pipeline, into sample-specific FASTQ files. These FASTQ files were then processed with the cellranger count pipeline where each sample was processed independently. First, cellranger count used STAR to align cDNA reads to the hg19 human reference transcriptome, which accompanied the Cell Ranger Single Cell Software Suite^43^. We note that, since the expression data is limited to the 3’ end of a gene and we used gene-level annotations, differences between reference versions, such as GRCh38, are unlikely to significantly alter conclusions. Aligned reads were filtered for valid cell barcodes and unique molecular identifiers (UMI) and observed cell barcodes were retained if they were 1-Hamming-distance away from an entry in a whitelist of known barcodes. UMIs were retained if they were not homopolymers and had a quality score > 10 (90% base accuracy). Cellranger count corrected mismatched barcodes if the base mismatch was due to sequencing error, determined by the quality of the mismatched base pair and the overall distribution of barcode counts. A UMI was corrected to another, more prolific UMI if it was 1-Hamming-distance away and it shared the same cell barcode and gene. Cellranger count examined the distribution of UMI counts for each unique cell barcode in the sample and selected cell barcodes with UMI counts that fell within the 99th percentile of the range defined by the estimated cell count value. The default estimated cell count value of 3,000 was used for this experiment. Counts that fell within an order of magnitude of the 99th percentile were also retained. The resulting analysis files for each sample were then aggregated using the cellranger aggr pipeline, which performed a between-sample normalisation step and merged all samples into one. Post-aggregation, the count data was processed and analysed using a comprehensive pipeline assembled and optimised in-house as described below.

To pre-process the mapped data, we constructed a cell quality matrix based on the following data types: library size (total mapped reads), the total number of genes detected, percent of reads mapped to mitochondrial genes, and percent of reads mapped to ribosomal genes (**Figure S13**). Cells that had any of the four parameter measurements that were greater than 3x median absolute deviation (MAD) of all cells were considered outliers and removed from subsequent analysis. In addition, we applied two thresholds to remove cells with mitochondrial reads above 20% or ribosomal reads above 50%. To exclude genes that were potentially detected from random noise, we removed genes that were detected in fewer than 1% of all cells. These quality control filters resulted in consistent total reads per individual and per pool in both fibroblasts and iPSCs (**Figure S14**). Before normalisation, abundantly expressed ribosomal genes and mitochondrial genes were discarded to minimise the influence of those genes in driving clustering and differential expression analysis.

### Demultiplexing

We adapted the Demuxlet method to our 10x scRNAseq data.^28^ The likelihood that a cell originated from a sample is the cumulative likelihood of single nucleotide polymorphism genotypes identified in each cell. We calculated posterior probability of a genotype g identified for a cell based on scRNA-seq data given the DNA data from the imputed BeadChip genotypes. Since the single cell SNP genotype data is sparse, to increase the coverage of SNPs called from scRNA-seq data that are in the SNP genotype data, we imputation SNP genotypes using the haplotype reference panel. We applied an ensemble approach using the outputs from pre-imputed genotype data, imputed genotype likelihood data, and impute genotype dosage data, increased the singlet probabilities from Demuxlet (**Figure S15**). The ensemble approach enabled the unique donor assignment of 90.6% of all cells, with high confidence to each sample, where demuxlet predicted no ambiguously assigned droplets. Of note, 100% of the cells before Demuxlet were identified in cellranger pipeline as a singlet. Demuxlet identified 90.6% of all cellranger singlet cells as ‘real’ single cells. Therefore, these cells were ascertained as singlets. To recover the cell assignment to the remaining 9.4% cell ranger singlets, predicted as doublets by Demuxlet, we utilised gene expression matrix to model cell doublets, using a simulation-based approach^44^. For each cell that was identified as both a singlet by demuxlet and the doublet expression simulation, was assigned to a donor based on the highest likelihood probability from demuxlet.

### Normalisation

Normalisation was conducted at four levels: between samples within a pool, between pools, between cells, and between clusters. The between-pool normalisation followed the subsampling strategy in the cellranger pipeline, where the reads, genes and cells were randomly subsampled following subsampling rates determined by the total read per sample and binomial distribution.^45^ Four pools were randomly multiplexed into one sequencing lane. For cell-to-cell normalisation, a cell-pooling strategy was applied to circumvent the zero-inflation issue, as described by Lun et al.^46^ Between pool normalisation followed Combat parametric empirical Bayesian strategy. To select the normalisation strategy, we compared results from using Combat, RUV and SCRAN methods by using k-BET batch-effect scores^32^. We found that a combination of SCRAN normalisation followed by Combat was superior in reducing batch effects compared to other methods, consistent with the results reported by Buttner and colleagues^32^. Prior to eQTL analysis, the mean expression of each gene per individual per cell subpopulation was computed and Z-transformed for eQTL mapping.

### Imputation and Quality control of genotype data

The 79 cell lines were genotyped by Infinium HumanCore-24 v1.1 BeadChip assay (Illumina). GenomeStudioTM V2.0 (Illumina) was used for SNP genotype calling of the BeadChip data (total 306,670 SNPs for one assay). The full genotype report files were reformatted into Plink map, fam, and lgen files and were then converted into variant calling format (vcf) using custom shell scripts and Plink2^47^. Plink2-converted files contained predicted reference and alternative alleles with no information for homozygous genotypes, which were fixed using the GenomeStudio report file and a custom script. For each sorted, indexed vcf file (separated by chromosomes), a strand fixing step was performed using bcf fixref function^48^. Prior to imputation, Eagle V.2.3.5 was used for haplotype phasing the strand-fixed genotype vcf files^49^. The phased data were imputed based on the 1000 genome phase 3 reference panel (2,535 samples) using the minimac3 program in the Michigan Imputation server^50^.

### Cell type classification and annotation

SingleR^30^ was used to map single cell transcriptomes against 713 reference transcriptomes. Then, we combined all cells from the fibroblasts and iPSCs pools separately. Using these two merged datasets, we normalised and clustered cells, ensuring the clustering was not affected by pool-specific data processing. We performed clustering using the SCORE method to identify subpopulations of cells^51^. Clustree^52^ was used to display the cluster stability at different resolutions (**Figure S16**). To visualise cell distributions, we used non-linear Uniform Manifold Approximation and Projection (UMAP) dimensionality reduction^53^. Cyclone^54^ was used to estimate cell cycle stages of each cell. Pseudo-trajectory analysis was carried out with slingshot^55^ using the UMAP cell projections.

### eQTL association analysis

To study specific regulation effects of genomic variance to gene expression, we performed statistical analysis of the association between genotypes of single nucleotide polymorphisms and single-cell gene expression for 79 fibroblast cell lines and 31 iPSC cell lines generated from the same individuals. We filtered for common SNPs (minor allele frequency > 0.05) that were within +/− 1 Mb of an expressed gene (detected in > 1% of the cells), resulting in 5,368,223 SNPs and 9,796 genes for the fibroblasts, and 4,508,778 and 10,899 genes for the iPSCs. SNP genotypes were recoded as 0, 1, or copies of the reference allele. eQTL mapping was performed for each subpopulation identified by the clustering analysis. Cis-eQTLs (SNP < 1 Mb) were detected using a linear model implemented in the MatrixEQTL R software with study-wide FDR lower than 5%^56^.

### Differential Expression

We used edgeR^57,58^ to identify differentially expressed genes between each cell type compared with the other cell types combined (i.e. each fibroblast type compared to the other five fibroblast types and each iPSC type compared to the other three iPSC types). Differentially expressed genes were detected using the gene-wise negative binomial generalised model with a quasi-likelihood test. Detection rate and pool batches were included as covariates following the recommendations of Soneson and Robinson^36^. Heatmaps and upset plots were generated using ComplexHeatmap^59^ in R. Heatmaps were created with scaled, normalised data.

### Independent eQTL analysis

Given an eGene that was significant in a pair of cell types (a and b), the top eSNPs from each cell type (S_a_ and S_b_) were tested for independency with relation to eGene expression. Accordingly, the top eSNP (S_b_) in cell type b was regressed from the linear model for the association of the top eSNP, S_a_, for cell type a with gene expression of the eGene (G_a_) in cell type a.

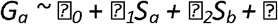

eSNPs were deemed independent if the association between S_a_ and G_a_ was significant following regression of S_b_ in the linear model.

### Interaction eQTL analysis

Given an eGene that was significant in at least two cell types, the eSNP with the largest difference between their beta allelic effects between any two clusters was used to test for cell type interaction. Two models were fit for gene expression G, with SNP S and cell type C. The first model (1) was a normal linear model and the second model (2) included an interaction term. An interaction was considered significant if an anova comparing the two models was significant.

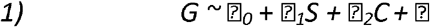

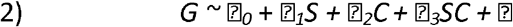

### eGene correlation

The expression of eGenes that were unique to a given cell type were correlated with their expression in the other cell types using a Pearson correlation test.

### eGene enrichment

eGenes from a specific cell type were tested for enrichment in the other cell types. eGenes were ranked based on the lowest P-value for each eGene. An expected distribution of mean rank scores were generated from 10,000 permutations of randomly selected genes (selecting the same number of genes as eGenes). The mean rank of the eGenes in the testing cell types were then tested for significance with a t-test.

### GTEx comparison

Gene Tissue Expression (GTEx)^33^ database version seven results were downloaded on 6 July, 2019. The cultured fibroblast cell eQTL were compared with the fibroblast cell type eQTL results to identify common and unique results.

## Tables

**Table 1:** Summary of fibroblast type *cis-eQTL*. The median number of cells per individuals, the number of significant eSNPs detected, the number of significant eGenes detected and the number of unique eGenes per cell type are enumerated.

