## Supporting figures and text for "Single cell eQTL analysis identifies cell type-specific genetic control of gene expression in fibroblasts and reprogrammed induced pluripotent stem cells"

**SUPPLEMENTARY FIGURES**

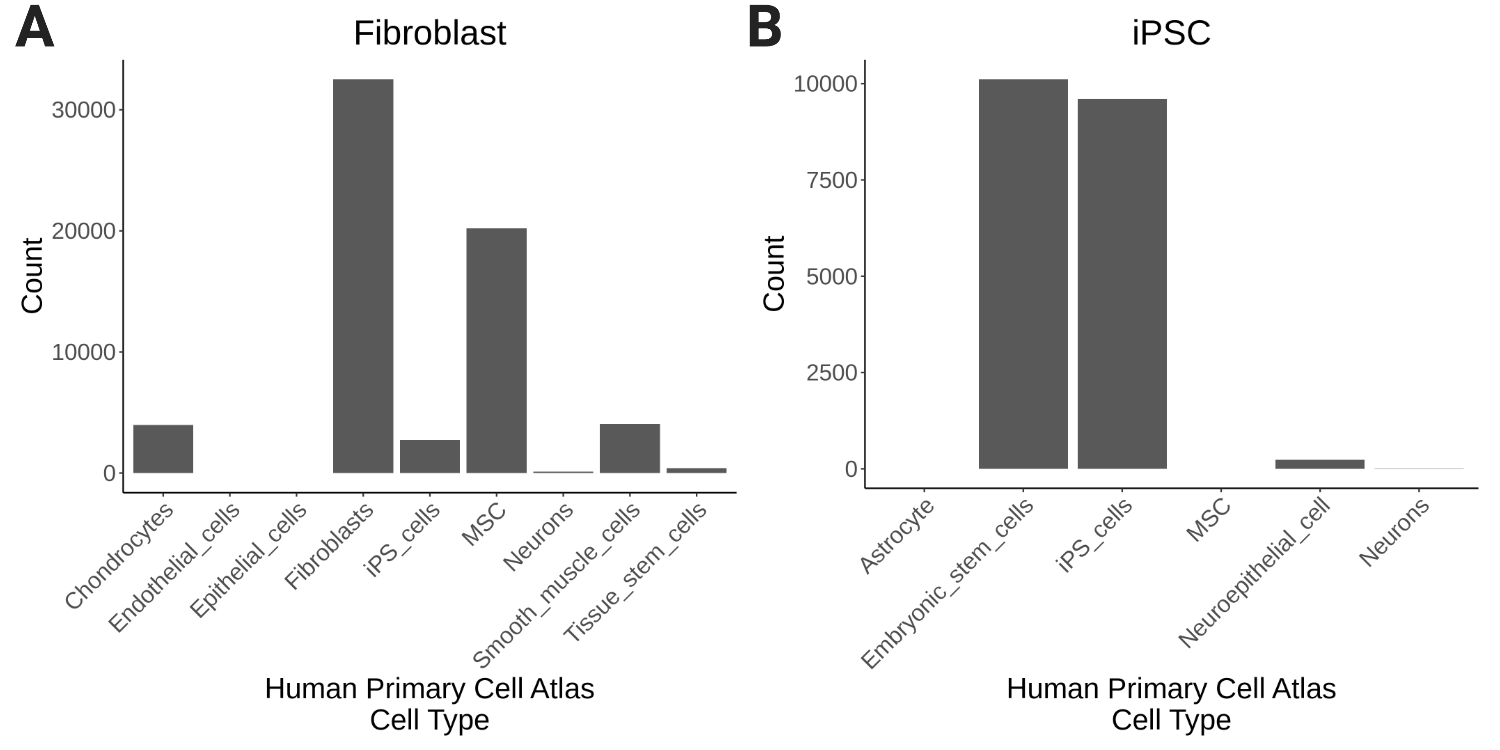

**Supplementary Figure S1: Map scRNA-seq Transcriptomes to Reference Datasets. A)** The transcriptional profiles of fibroblast cells were mapped to 713 reference transcriptomes available from the human primary cell atlas. The majority of these cells were mapped to the Fibroblast reference dataset. **B**) iPSC scRNA-seq transcriptional profiles were mapped to 713 reference transcriptomes from the human primary cell atlas. The vast majority of iPSC single cell transcriptional profiles were mapped to stem cells (embryonic or induced pluripotent stem cells).

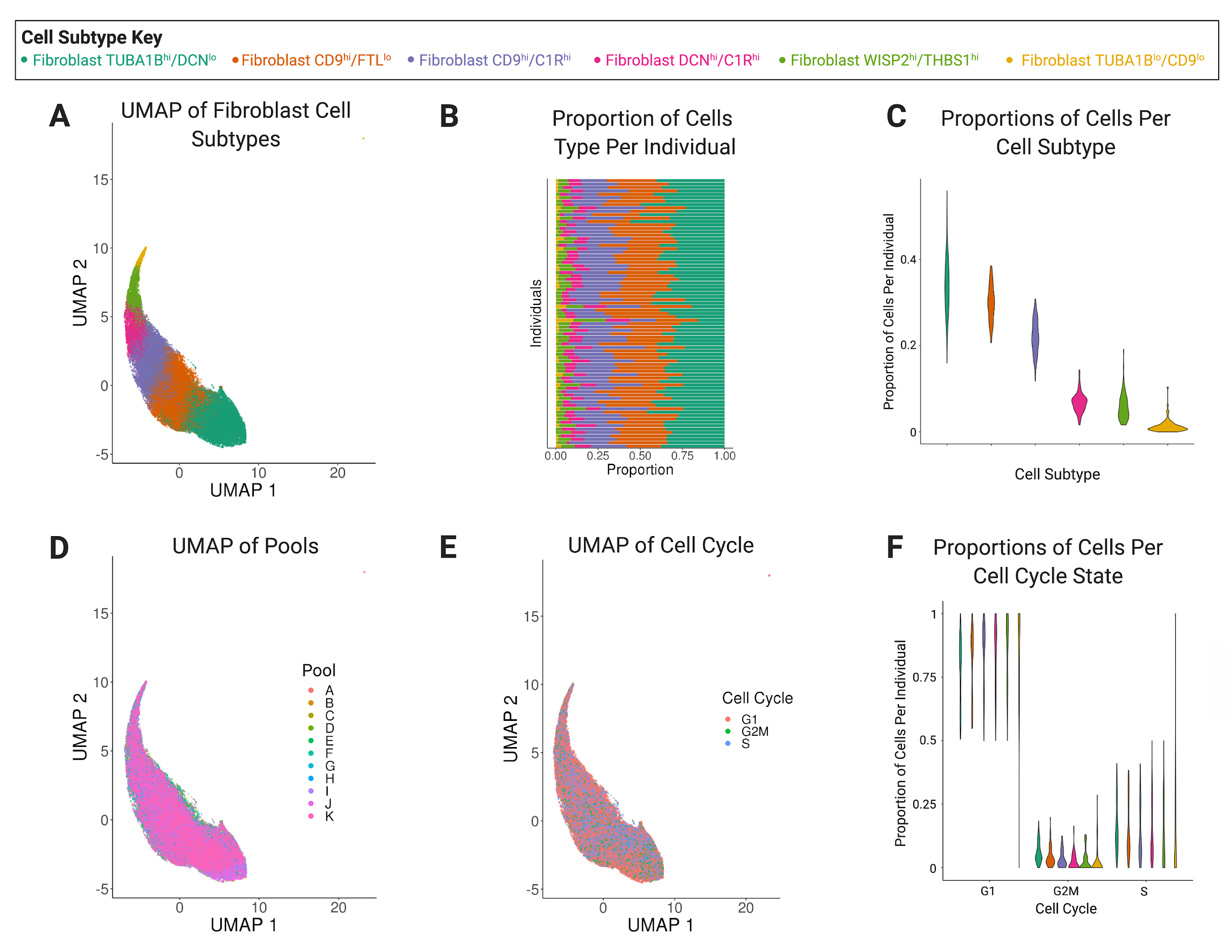
**Supplementary Figure S2: Quality Control Metrics of Fibroblast Subtypes. A)** Six subtypes of fibroblasts were identified. **B-C)** The proportions of each fibroblast subtype were consistent across individuals. **D)** The 10x capture pools were evenly distributed across the six fibroblast subtypes. **E-F)** Cell cycle states were also evenly distributed across fibroblast subtypes.

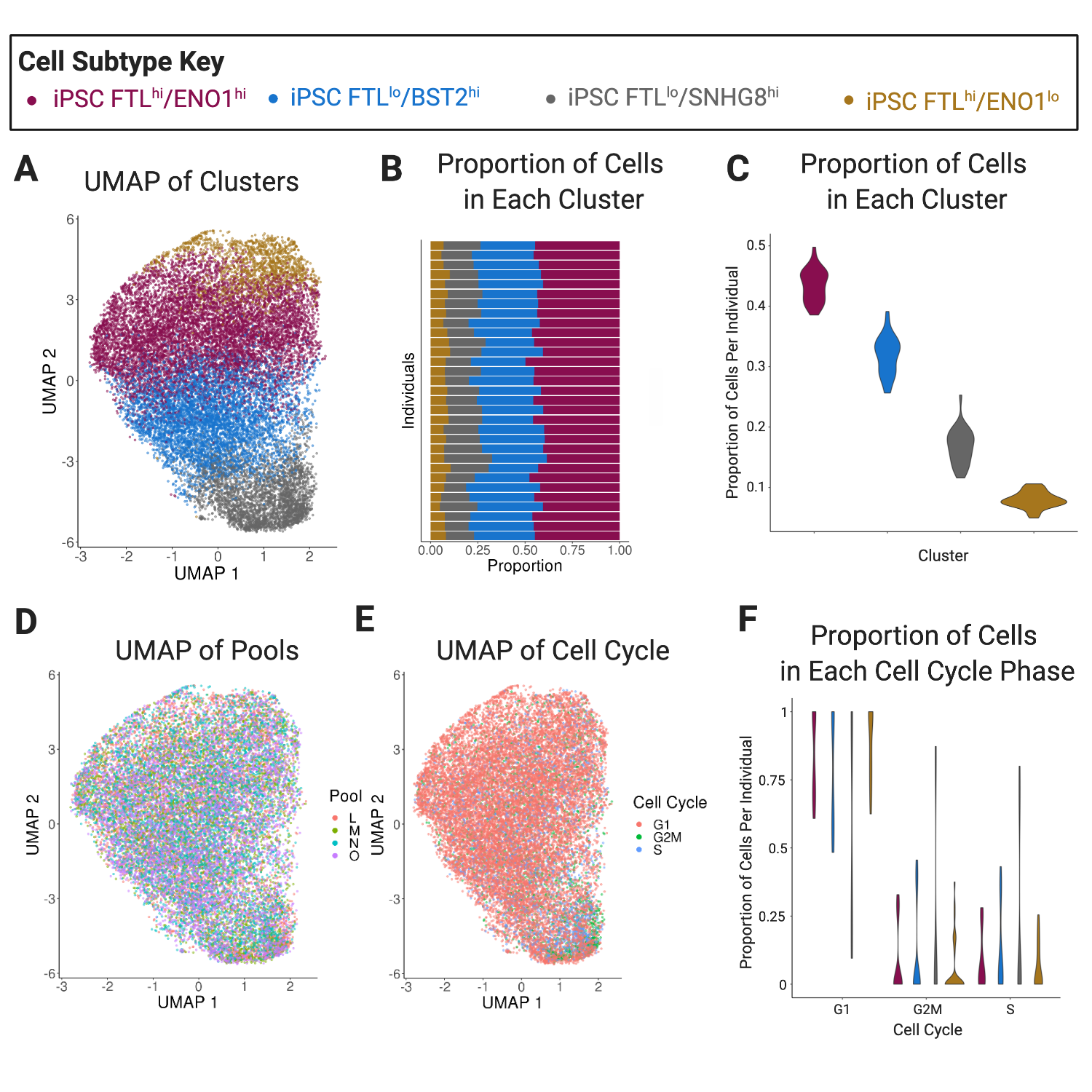

**Supplementary Figure S3: Quality Control Metrics of Fibroblast Subtypes. A)** Four subtypes of induced pluripotent stem cells (iPSCs) were identified. **B-C)** The proportions of each iPSC subtype were consistent across individuals. **D)** The 10x capture pools were evenly distributed across the six iPSC subtypes. **E-F)** Cell cycle states were also evenly distributed across iPSC subtypes.

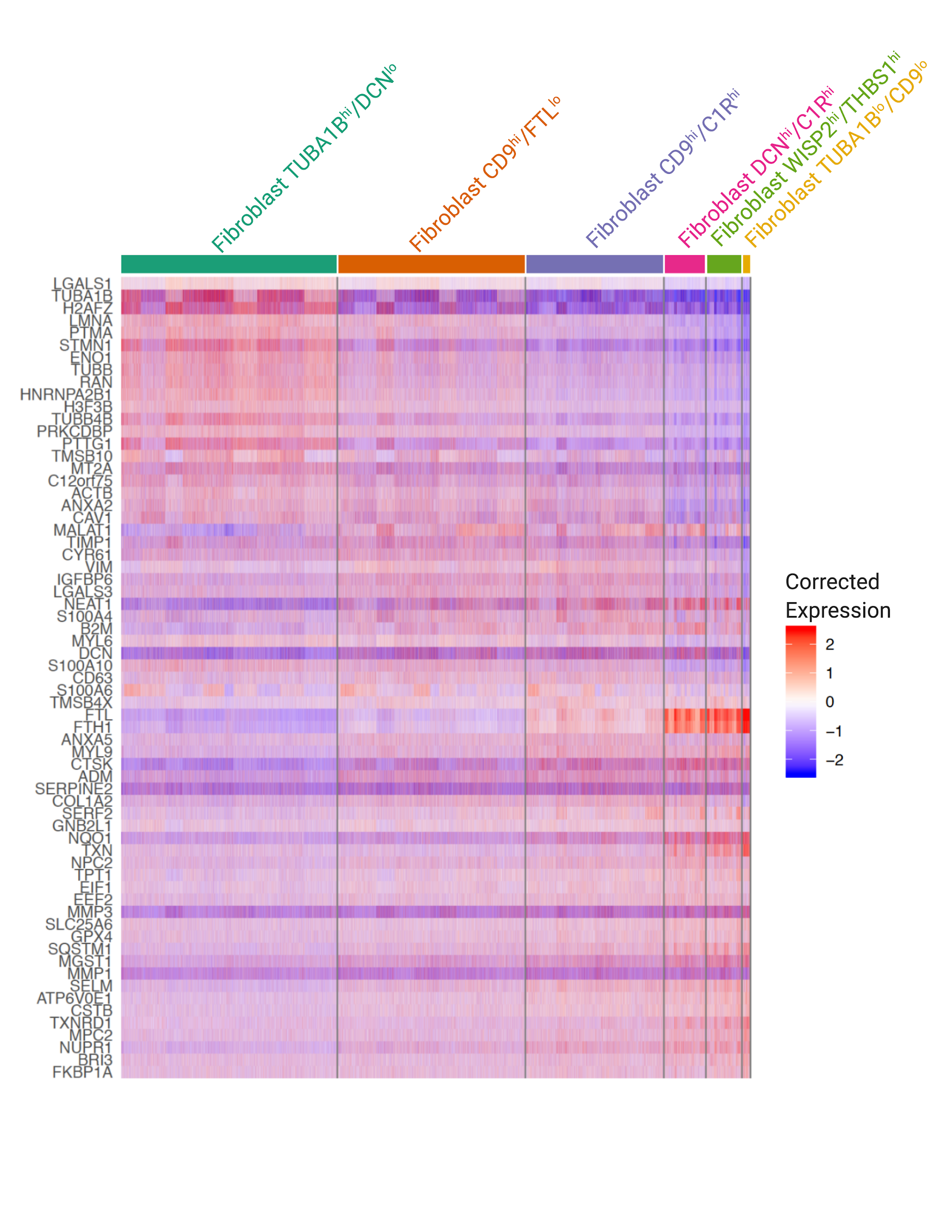

**Supplementary Figure S4: Top 20 differentially expressed genes across the six fibroblast subtypes.** Expression of top 20 differentially expressed genes in each cell across the six fibroblast subtypes.

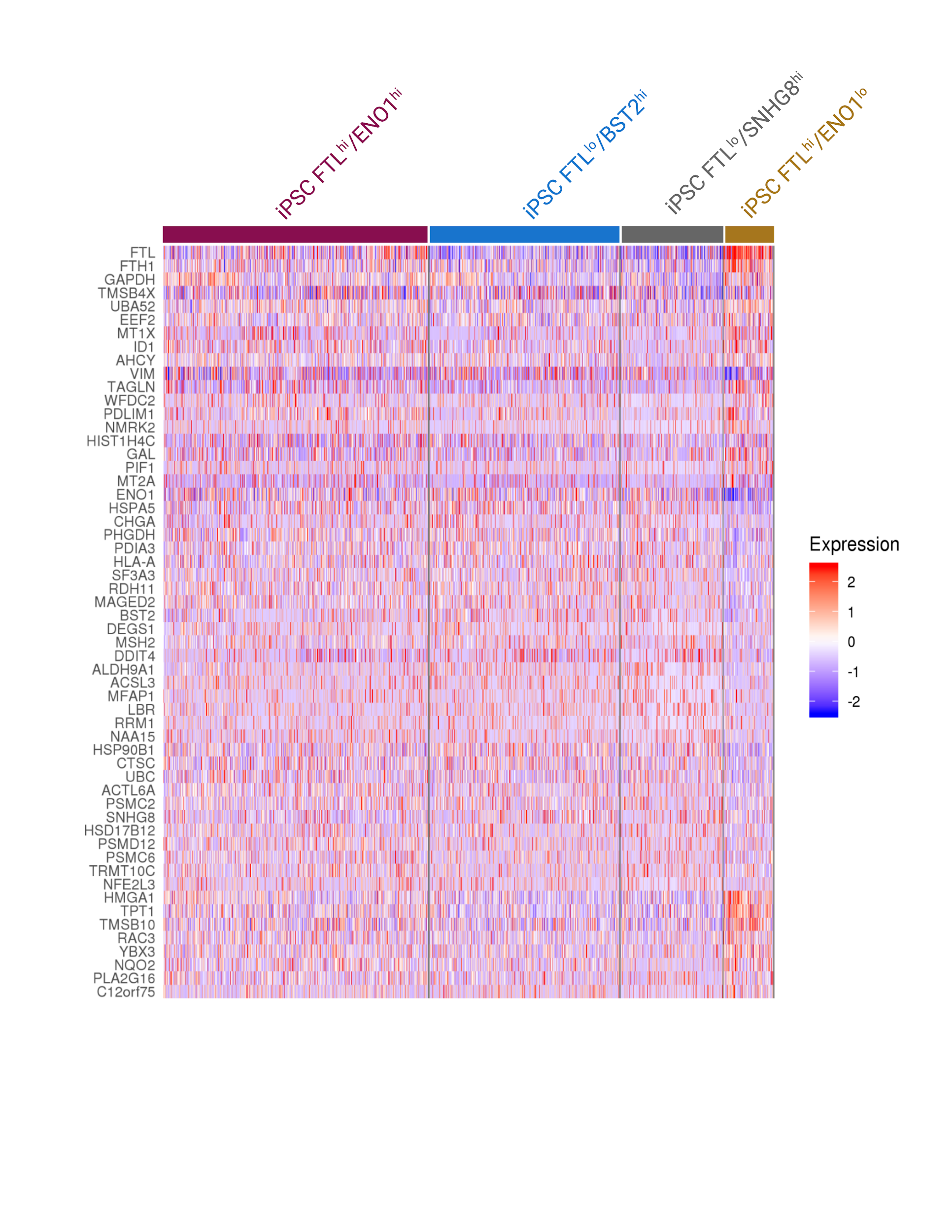

**Supplementary Figure S5: Top 20 differentially expressed genes across the four iPSC subtypes.** Expression of top 20 differentially expressed genes in each cell across the four iPSC subtypes.

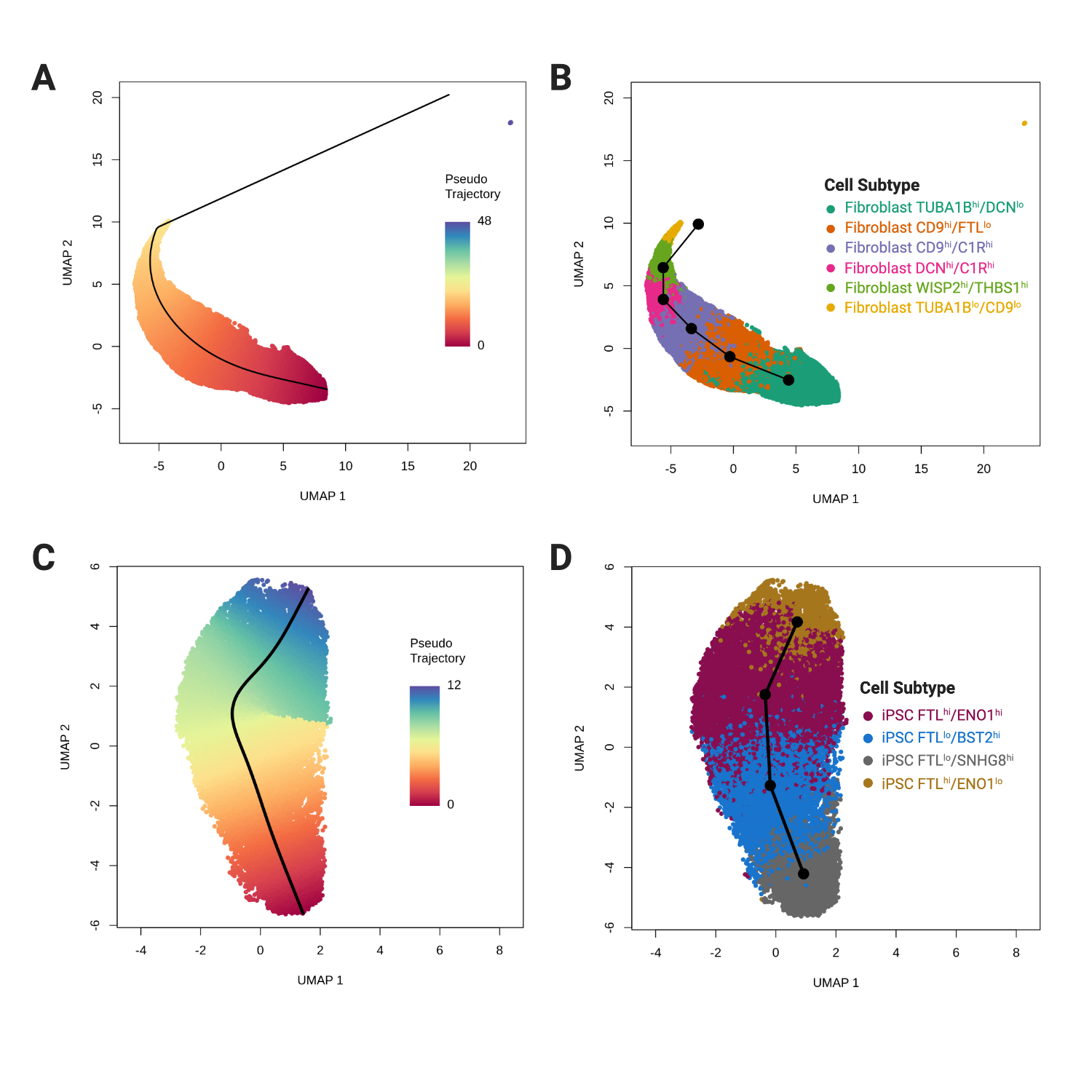

**Supplementary Figure S6: Fibroblast and iPSC Pseudotrajectories. A)** Pseudotrajectory across fibroblast single cells colored by the slingshot pseudotrajectory values B) Fibroblast subtype pseudotrajectory colored by the fibroblast subtype. C) Pseudotrajectory across iPSC single cells colored by slingshot pseudotrajectory values. D) iPSC subtype pseudotrajectory colored by the iPSC subtypes.

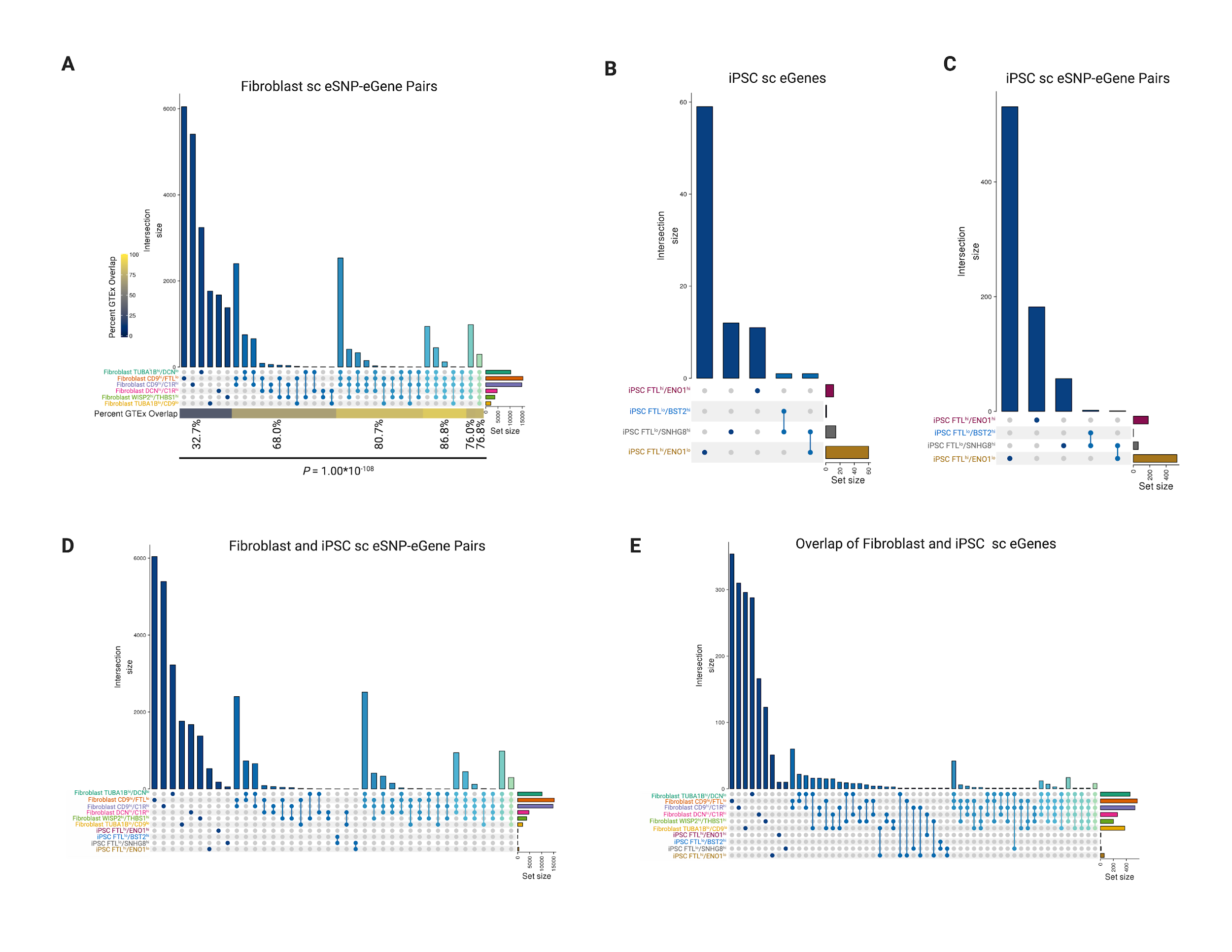

**Supplementary Figure S7: Overlap of sc eQTLs in Fibroblasts and iPSCs.** **A)** The majority of eSNP-eGene pairs in fibroblast subpopulations are unique to a given subpopulation. Further, the percent of those sc eSNP-eGene pairs that were detected in GTEx transformed fibroblasts increased with increasing numbers of fibroblast subtypes that they were detected in (P = 1E-108, Cochran-Armitage Test). **B)** The majority of iPSC eGenes were unique to an iPSC subtype and none were significant in more than two subtypes. **C)** In addition, the majority of iPSC eSNP-eGene pairs were also unique to a single iPSC subtype. **D)** No eSNP-eGene pair was significant in both fibroblast and iPSC subtypes. **E)** However, some eGenes were significant in both fibroblast and iPSC subtypes.

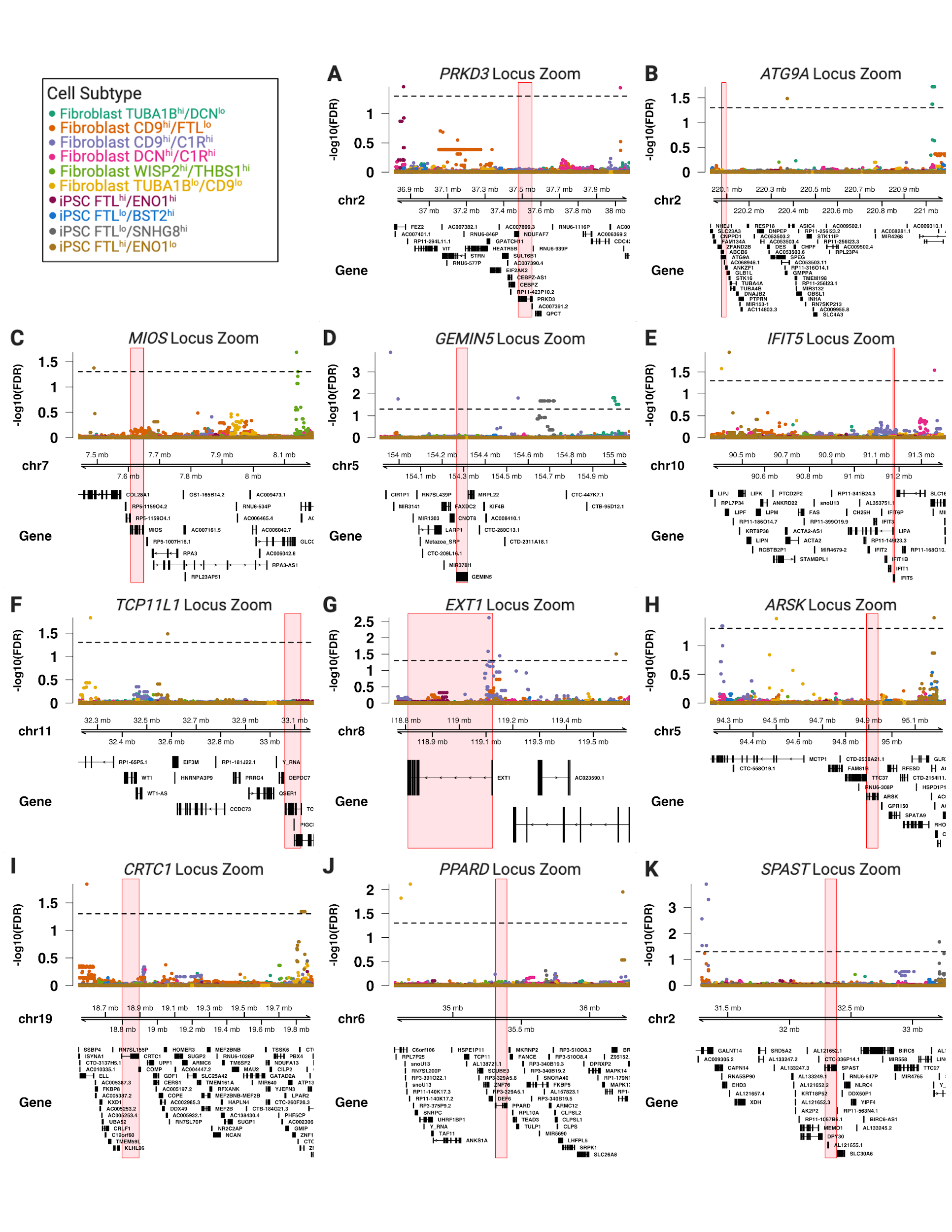

**Supplementary Figure S8: eGenes that were Significant in both iPSC and Fibroblast Subtypes.** Eleven eGenes were significant in at least one iPSC and one fibroblast subtype. Those included *PRKD3* (**A**), *ATG9A* (**B**), *MIOS* (**C**), *GEMIN5* (**D**), *IFIT5* (**E**), *TCP11L1* (**F**), *EXT1* (**G**), *ARSK* (**H**), *CRTC1* (**I**), *PPARD* (**J**) and *SPAST* (**K**). In all cases, the eSNPs for the fibroblast and iPSC subtypes were either on opposite sides of the gene or separated by at least 400,000 base pairs.

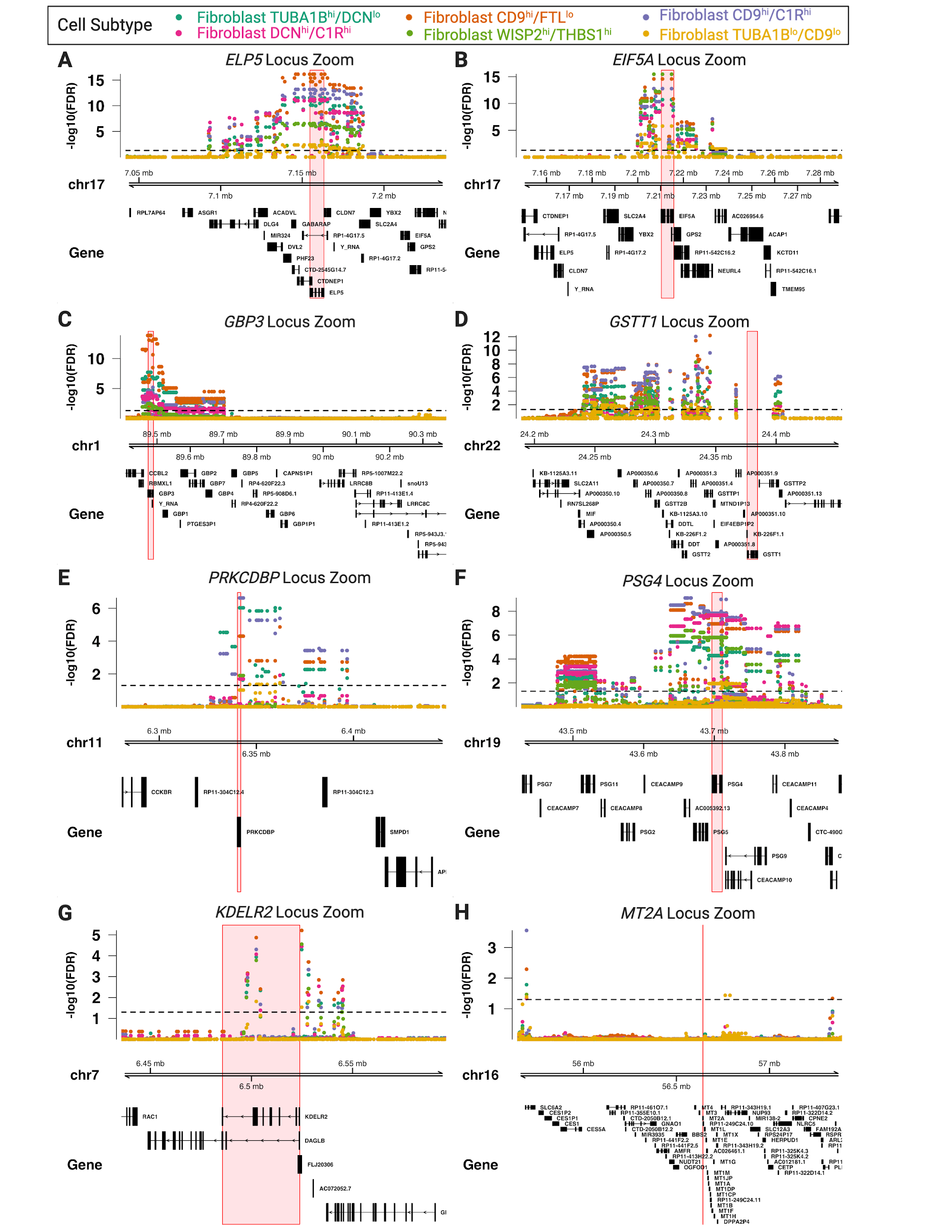
**Supplementary Figure S9: eGenes that were significant in all six fibroblast subtypes.** Eight eGenes were significant in all six fibroblast subtypes. Those included *ELP5* (**A**), *EIF5A* (**B**), *GBP3* (**C**), *GSTT1* (**D**), *PRKCDBP* (**E**), *PSG4* (**F**), *KDELR2* (**G**) and *MTA2* (**H**).

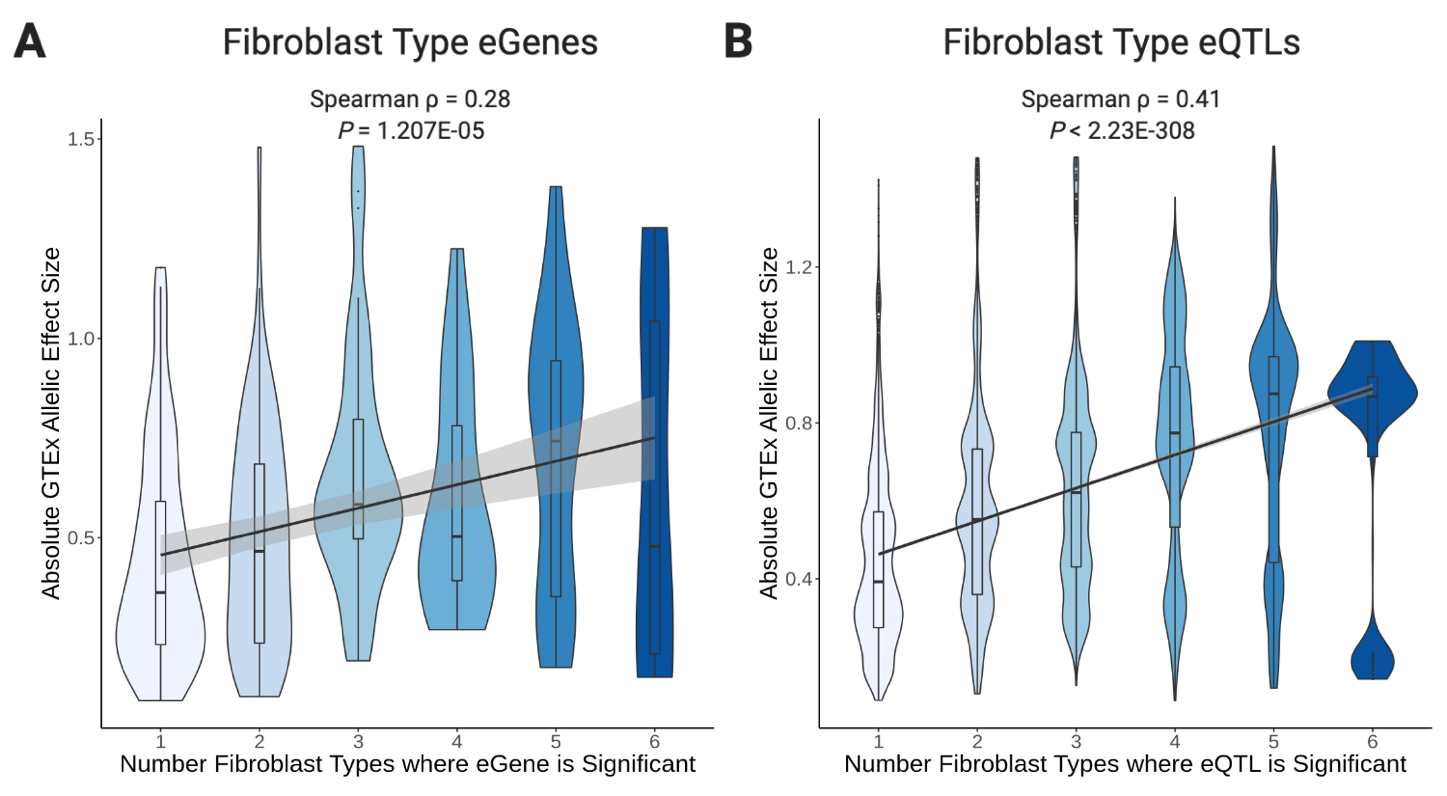

**Supplementary Figure S10: Absolute GTEx Cultured Fibroblast Allelic Effect Sizes by Number of Significant Fibroblast Types.** **A)** The absolute allelic effect size from GTEx cultured fibroblasts is significantly correlated with the number of fibroblast types where that eGene was significant. The absolute allelic effect size in GTEx cultured fibroblasts was smallest for those eGenes that were significant in just one fibroblast type. **B)** The absolute allelic effect size from GTEx cultured fibroblasts is significantly correlated with the number of fibroblast types where that eQTL was significant. The absolute allelic effect size in GTEx cultured fibroblasts was smallest for those eQTLs that were significant in just one fibroblast type.

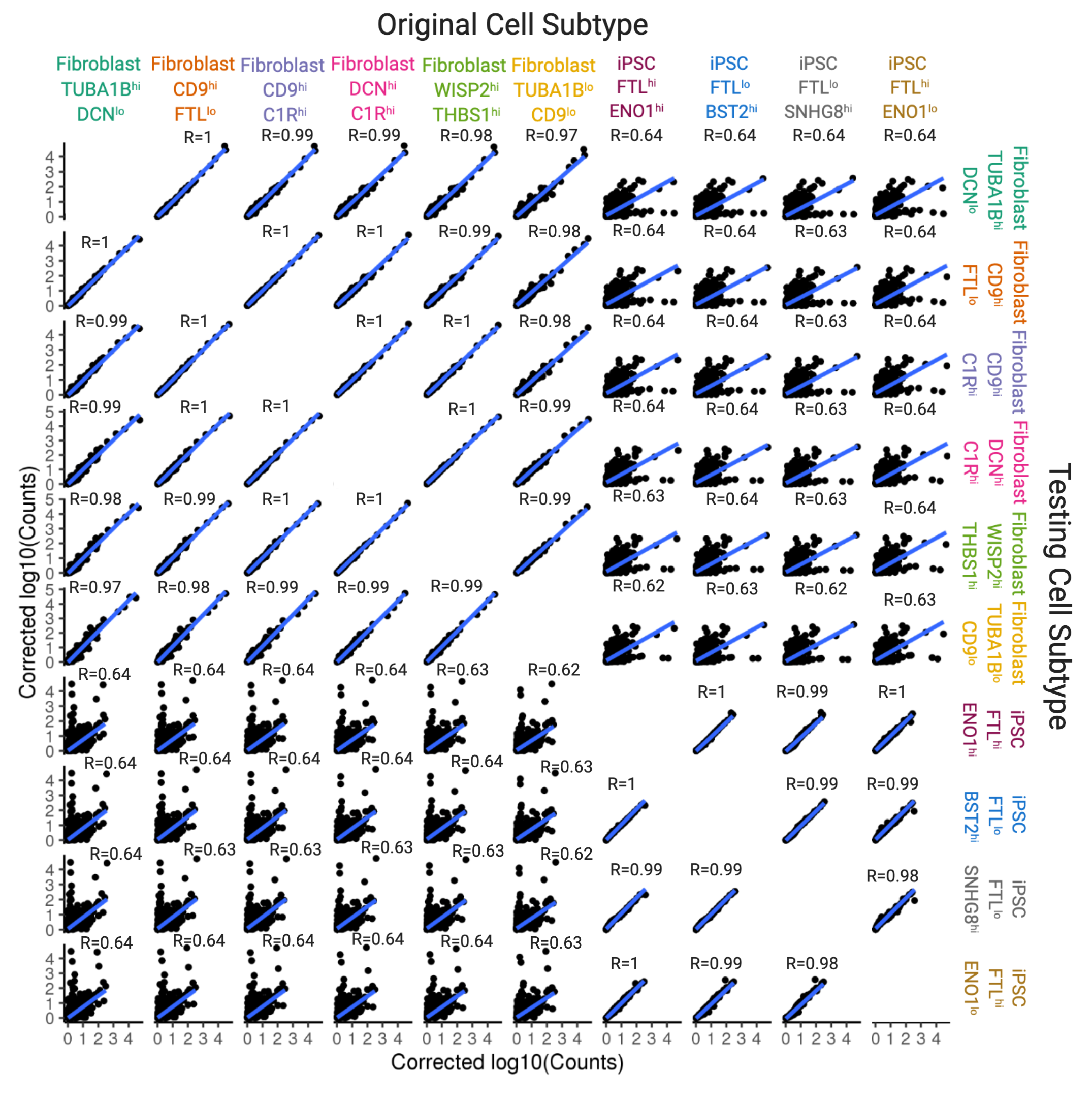

**Supplementary Figure S11: Correlation of eGene Expression.** eGene expression from the one cell subtype (“Original Cell Subtype”) were correlated for expression in the other cell subtypes (“Testing Cell Subtype”). Pearson correlation was used to test the linear relationship of the “Original Cell Subtype” eGene expression with expression in the “Testing Cell Subtype”.

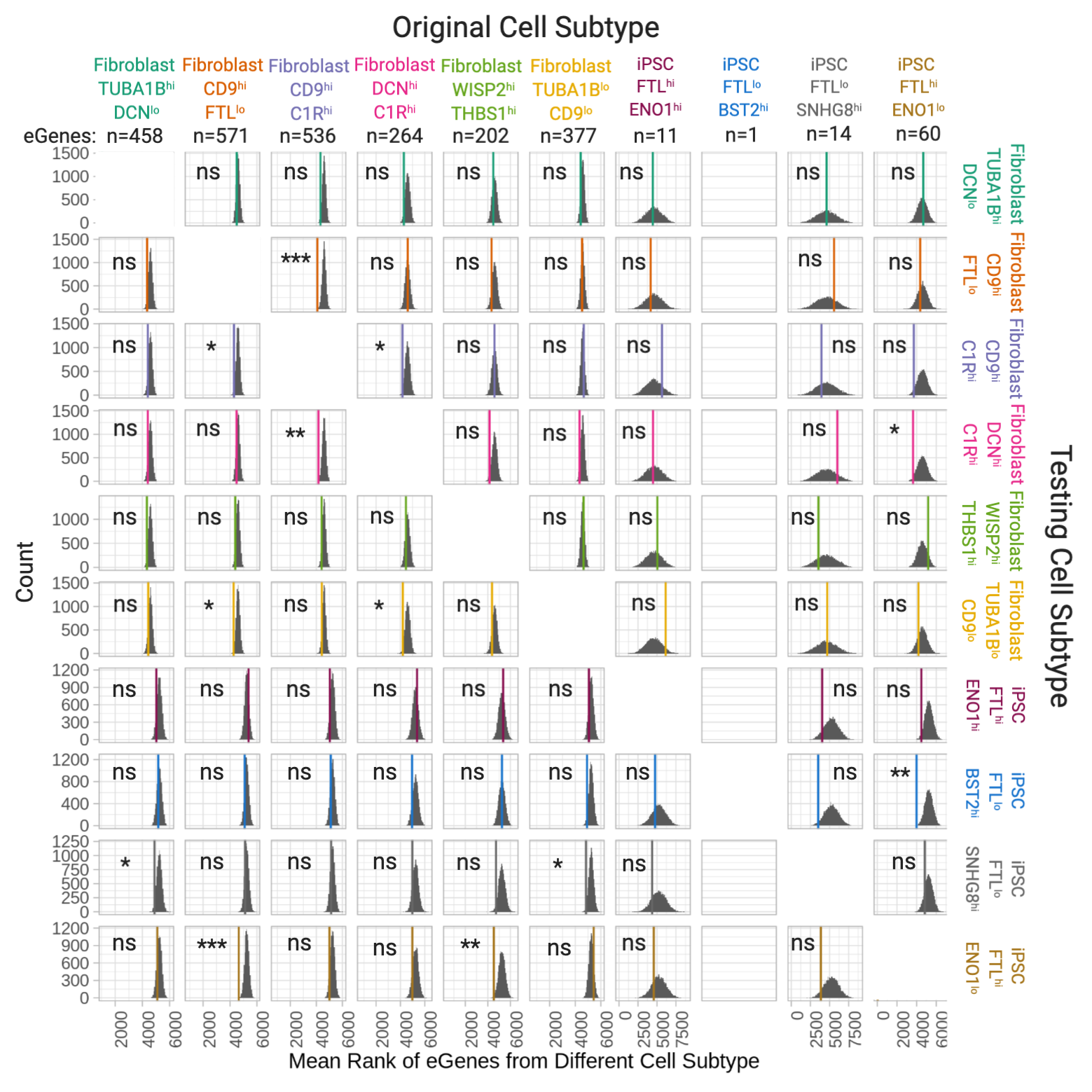

**Supplementary Figure S12: Enrichment of Unique Significant eGenes in other Subtypes.**

Significant eGenes from each cell subtype were tested for enrichment in other cell subtypes. The grey densities represented 10,000 permutations of mean rank of randomly-selected eGenes, the colored line is the mean rank of the eGenes from the original cell subtype in the testing cell subtype. The iPSC FTL^lo^/BST2^hi^ cell subtype had too few unique eGenes (1) to test for enrichment in other cell subtypes. A student’s t-test was used to test if the mean rank of the eGenes was significantly different from the genes randomly selected from the testing cell subtype eGene ranked list. **P* < 0.05; ***P* < 0.01; ****P* < 0.001; ns=non-significant.

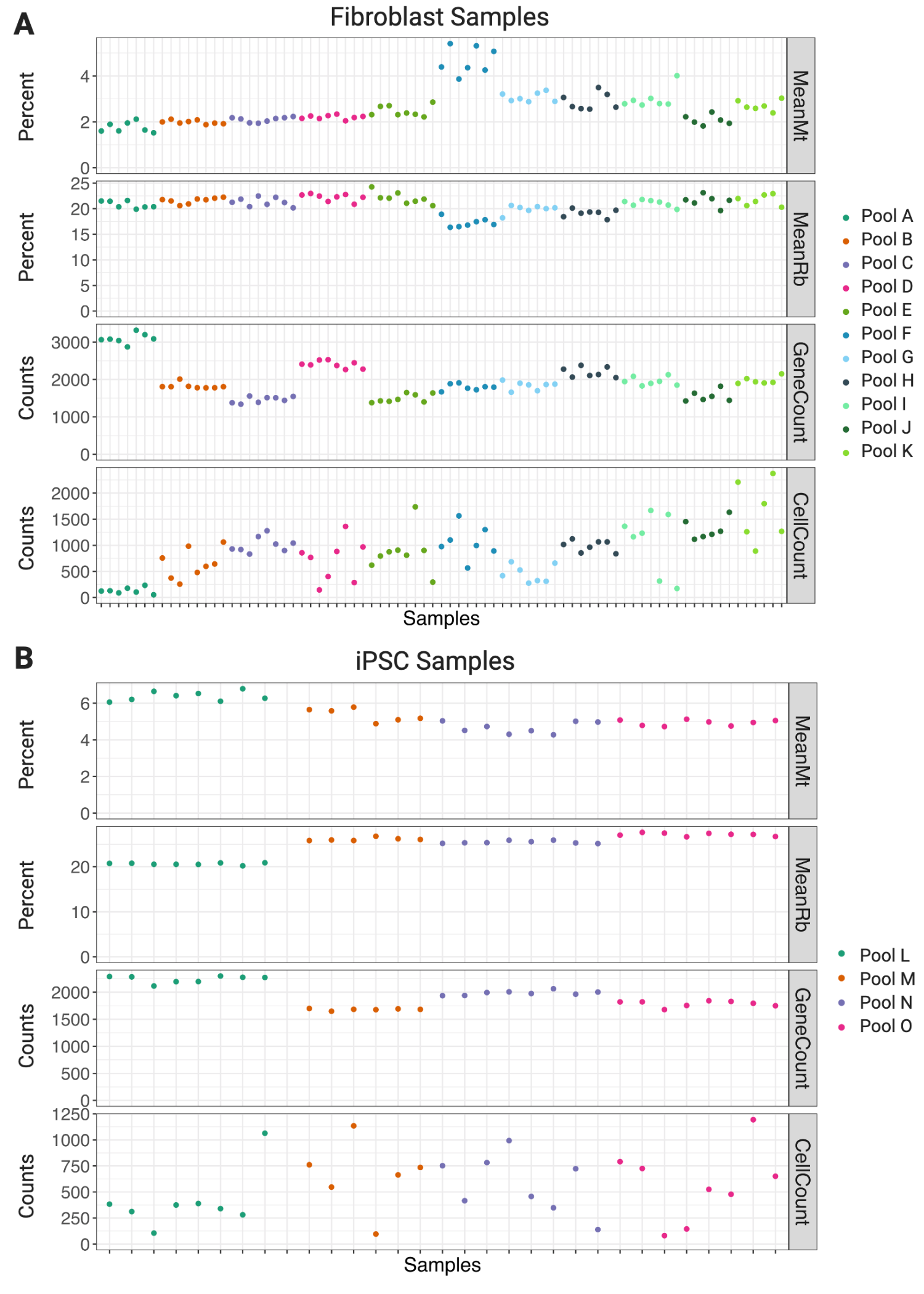

**Supplementary Figure S13: Average Cell Quality Measures by Sample.** Mean percent of unique reads that map to mitochondrial genes (MeanMt), mean percent of unique reads that map to ribosomal genes (MeanRb), mean number of genes detected per cell and mean number of cells per sample before quality control processing in fibroblast (**A**) and iPSC (**B**) samples.

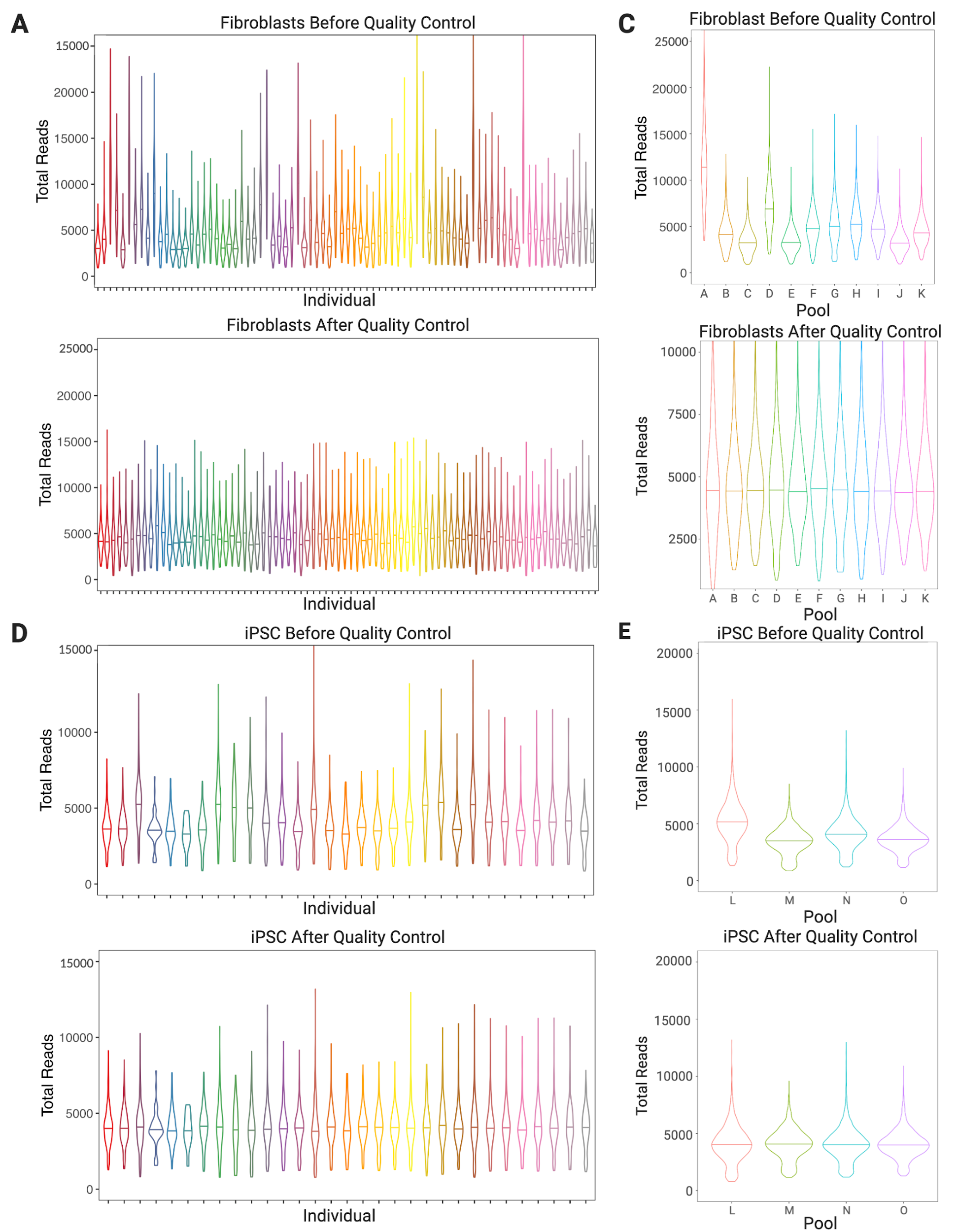

**Supplementary Figure S14: Total Reads per Cell Before and After Quality Control Filtering.** Quality control filtering resulted in consistent distributions of the total number of reads per cell across individuals in fibroblasts (**A**) and iPSC (**B**) as well as across pools in fibroblasts (**C**) and iPSC (**D**).

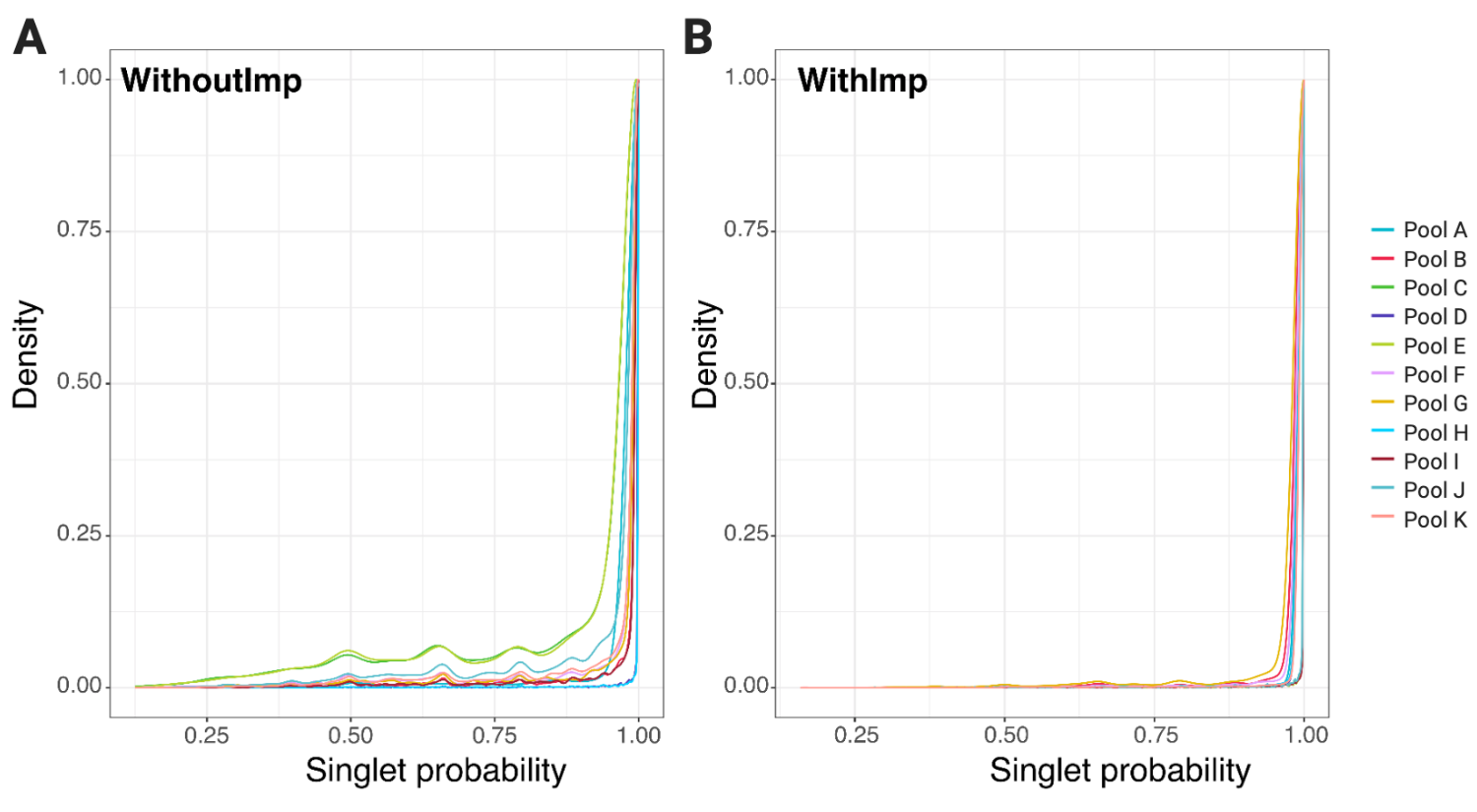

**Supplementary Figure S15: Distributions of Singlet Probability Without and With Imputed SNP genotypes. A**) Singlet probability distributions were were lower when demuxlet was applied without imputed SNPs (WithoutImp) than **B**) when demuxlet was applied with imputed SNP genotypes (WithImp).

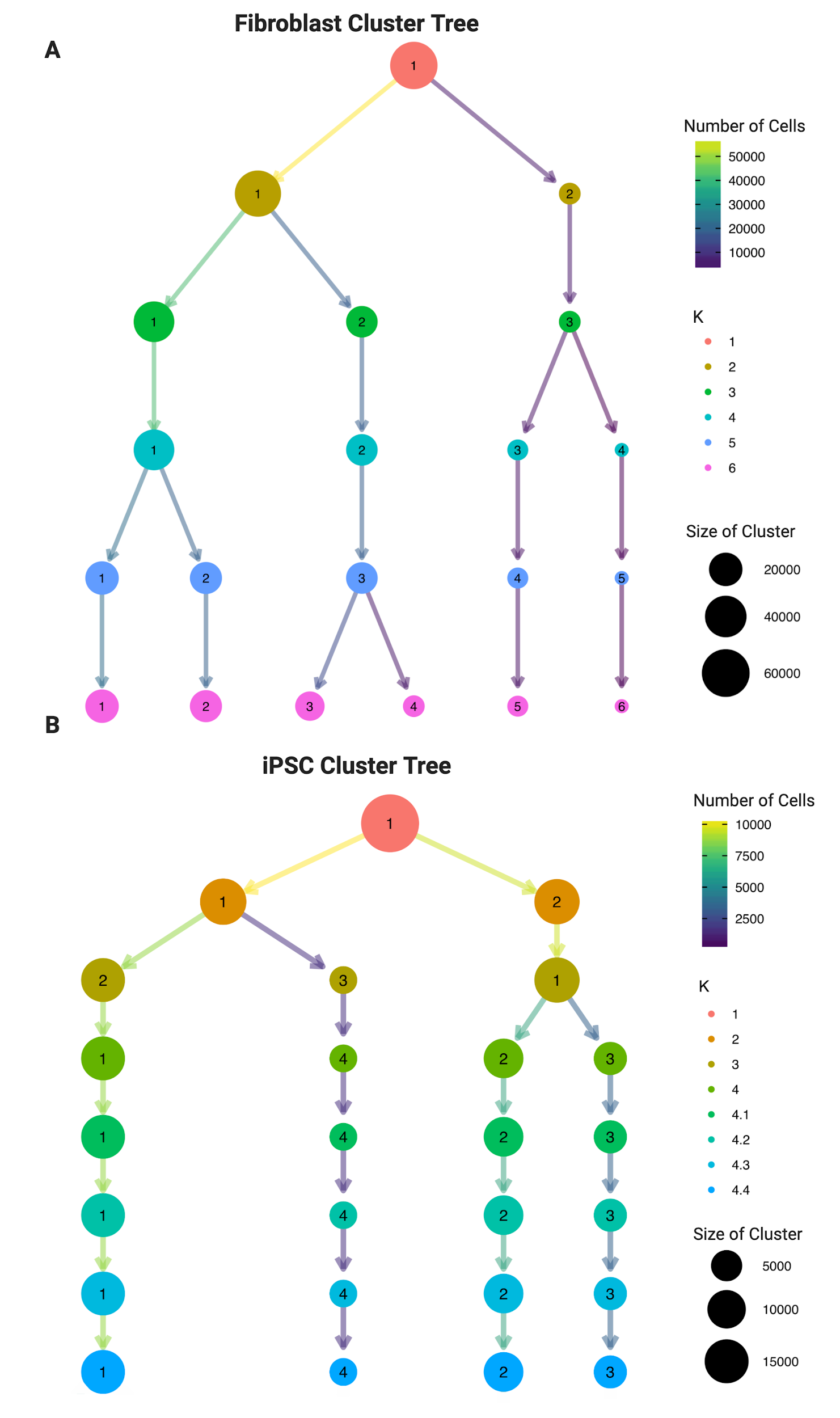

**Supplementary Figure S16: Cluster Tree of fibroblast (A) and iPSC single cell subtypes (B).** Four subtypes of induced pluripotent stem cells (iPSCs) were identified using SCORE clustering.
